# Large extracellular vesicles derived from red blood cells in coronary artery disease patients with anemia promote endothelial dysfunction

**DOI:** 10.1101/2025.03.10.642191

**Authors:** Isabella Solga, Vithya Yogathasan, Patricia Wischmann, Tin Yau Pang, Leon Götzmann, Patricia Kleimann, Sebastian Temme, Lina Hofer, Anja Stefanski, Alexander Lang, Georg Nickenig, Christian Jung, Norbert Gerdes, Mohammed Rabiul Hosen, Malte Kelm, Ramesh Chennupati

## Abstract

**Background and purpose:** Endothelial dysfunction (ED) is a hallmark of cardiovascular disease (CVD). We recently showed that anemia is associated with worsening of endothelial function after acute myocardial infarction (AMI). Extracellular vesicles (EVs) are efficient communicators between cells and can functionally contribute to different CVD, including, AMI. However, their specific role of EVs in stable coronary artery disease (CAD)-associated with anemia, particularly their contribution to ED, has not yet been investigated systematically.

**Experimental approach:** Red blood cell-derived EVs (REVs) and plasma-derived EVs (PLEVs) from all blood cells and endothelium were isolated from patients with stable CAD. The isolated large REVs and PLEVs were characterized using dynamic light scattering (DLS), nanoparticle tracking analysis (NTA), transmission electron microscopy (TEM), and Western blotting. Uptake assays were performed by co-incubating with fluorescently-labeled REVs and PLEVs with human umbilical vein endothelial cells (ECs). Nitric oxide (NO) consumption ability of REVs was analyzed using a chemiluminescence detector (CLD). After co-incubation of aortic rings explanted from wild-type (WT) mice with REVs and PLEVs from anemic and non-anemic CAD patients, endothelial function was assessed using a wire myograph system. To investigate differences in the content of REVs and PLEVs between anemic and non-anemic CAD patients, proteomic analysis was performed.

**Key results:** DLS analysis showed that both REVs and PLEVs were within the size distribution range of 100-1000 nm. NTA analysis revealed increased release of REVs in anemic patients compared to non-anemic patients. Co-incubation of labeled REVs and PLEVs with ECs demonstrated their uptake by ECs *in vitro* which was similar between anemic patients compared to non-anemic patients. REVs from anemic patients showed increased NO consumption compared to those from non-anemic patients. Aortic rings co-incubated with REVs from anemic patients showed attenuated endothelial NO-dependent relaxation responses compared to non-anemic patients. Proteomics analysis of REVs from anemic patients revealed numerous differentially expressed proteins, including decreased abundance of antioxidant proteins such as catalase 1 (CAT1), superoxide dismutase 1 (SOD1) and increased oxidative stress-promoting myeloperoxidase (MPO). Co-incubation of ECs with REVs from anemic patients demonstrated increased ROS production.

**Conclusion:** Anemia is associated with increased release of REVs and enhanced NO consumption, which promotes ED. This is further exacerbated by an altered redox balance and increased ROS production, implicating therapeutic importance in anemic patients with CAD.

**Graphical Abstract:** Anemia is associated with an increased release of RBC-derived large extracellular vesicles (REVs), which are taken up by endothelial cells (ECs). Anemic REVs show enhanced nitric oxide (NO) consumption, contributing to NO dysregulation in ECs. Additionally, REVs carry various redox enzymes, including the oxidative stress-promoting enzyme myeloperoxidase (MPO), as well as antioxidant enzymes such as superoxide dismutase (SOD) and catalase (CAT). An imbalance in these redox enzymes leads to increased oxidative stress and endothelial nitric oxide synthase (eNOS) uncoupling, resulting in impaired NO-mediated relaxation responses and subsequent endothelial dysfunction (ED).

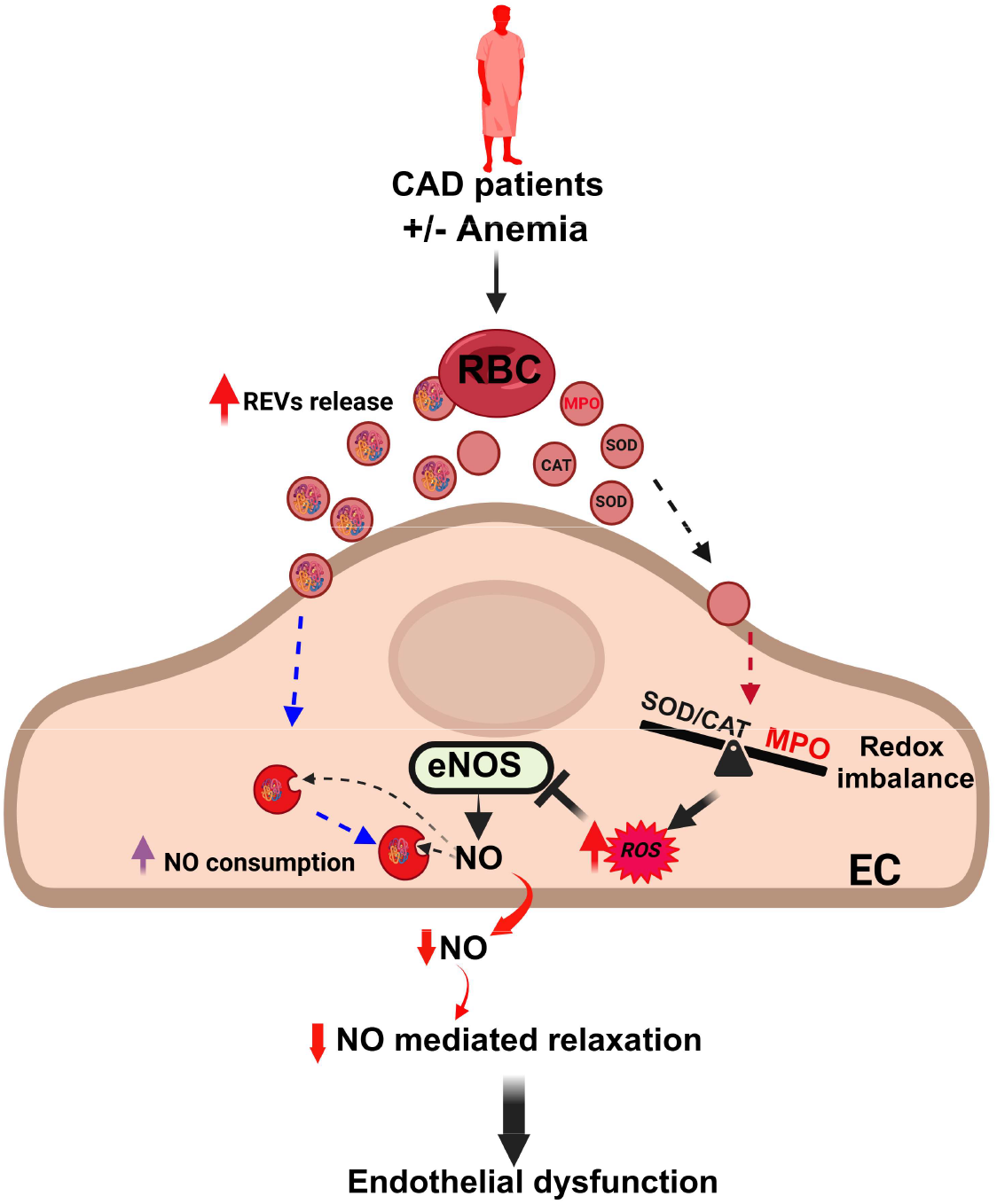

## Introduction

Extracellular vesicles (EVs) are released by different cells in the vascular system and play a crucial role in intercellular communication [1]. In recent years, the study of EVs has gained significant attention due to their involvement in various physiological and pathological conditions in the cardiovascular system [2]. EVs are heterogeneous membranous particles enclosed by a lipid bilayer [3]. Based on their origin and size, they are typically classified into three distinct subtypes [4]. Small particles, known as small EVs (formerly known as exosomes, 30-100 nm), large EVs or (formerly known microvesicles, 100-1000 nm) and apoptotic bodies (1-5 *μ*m) [4]. Large EVs are secreted by direct outward budding or shedding from the plasma membrane of the parental cell [5]. The formation of EVs is triggered by different stimuli such as shear stress, cell injury, cytokines, adenosine triphosphate (ATP) depletion and calcium influx [6].

The content of EVs is highly variable and includes proteins such as surface receptors, signalling proteins, transcription factors, enzymes but also carry lipids and nucleic acids such as miRNA, mRNA and DNA [7]. These bioactive cargos are important regulators of cell function via EV-mediated horizontal transfer in cellular crosstalk [8-10]. When released from donor cells, EVs interact with recipient cells, triggering intercellular signalling and altering molecular processes, potentially affecting their physiological or pathological states [11]. Previous studies showed elevated EVs in patients with cardiovascular risk factors (e.g., diabetes mellitus, hypertension, and hypercholesterolemia) as well as in those with cardiovascular diseases (CVD) or myocardial disorders [12, 13]. Furthermore, EVs are also considered clinical markers for inflammation and tissue/organ damage, thus potentially contributing to therapeutic approaches such as cardiovascular regeneration and protection [14].

Red blood cells (RBCs) are primarily known for their function of transport of gases, but they are also involved in the maintenance of vascular homeostasis through the regulation of nitric oxide (NO) pool and the regulation of redox balance [15, 16]. In our recent studies, we demonstrated that anemia is associated with RBC and endothelial dysfunction (ED) [17, 18]. In the same study we showed that chronic anemia is associated with inflammation and increased ROS formation that plays a crucial role in anemia associated ED [18]. Additionally, co-incubation of anemic RBCs from both mice and humans with murine aortic segments revealed impaired endothelial function, suggesting that anemic RBCs play a crucial role in mediating ED in anemia. However, the exact mechanisms by which anemic RBCs contribute to ED is not fully understood. RBC-derived EVs (REVs) have been shown to play a crucial role in various pathological conditions, including sickle cell anemia, thrombosis, CVDs, diabetes, cancer, and inflammation [6, 19, 20]. They contain a wide range of different compounds, including enzymes involved in redox homeostasis such as glutathione S transferase (GST), thioredoxin and peroxiredoxin but also myeloperoxidase (MPO), cholesterol and haemoglobin (Hb). These factors can affect vascular inflammation, reactive oxygen species (ROS) production, endothelial damage, and coronary heart disease. Additionally, the REVs are known to be potent NO scavengers, which reduce NO bioavailability and influence vasoregulation [6, 20]. The role of REVs in mediating ED is not clearly understood. We hypothesize that REVs play a central role in the communication between RBCs and endothelial cells in anemia, contributing to the development and progression of ED.

In this study, we investigated the role of anemic RBC-derived extracellular vesicles (REVs) in mediating endothelial dysfunction (ED) using samples from stable coronary artery disease (CAD) patients, aiming to uncover the impact of intercellular communication in these patient cohorts. Furthermore, from the same patients, we will investigate the role of anemic plasma-derived extracellular vesicles (PLEVs) in ED, which are derived from various cell types including RBCs, endothelial cells, platelets and other blood cells. To our knowledge, this is the first study to explore the specific role of anemic REVs and PLEVs in ED associated with anemia in CAD patients.

## Materials and Methods

### Collection of blood from patients

Blood samples from anemic and non-anemic patients were collected in EDTA tubes. Male patients were classified as anemic if their hemoglobin (Hb) levels were below 13.0 g/dL, while female patients were considered anemic if their Hb levels were below 12.0 g/dL (according to WHO guidelines). All patients included in this study had given their written consent and were recruited from the Department of Cardiology, Pulmonology, and Angiology at Düsseldorf University Hospital. The approval numbers for human blood samples collection are 5481R, 2018-14, and 2018-47, as approved by the ethics committee of Düsseldorf University Hospital.

### Mice

All animal procedures used in the study were approved and performed in accordance with the ARRIVE (Animal Research: Reporting of *In Vivo* Experiments) II guidelines and authorized by LANUV (North Rhine-Westphalia State Agency for Nature, Environment and Consumer Protection) in compliance with the European Convention for the Protection of Vertebrate Animals used for Experimental and other Scientific Purposes. The approval numbers for the animal experiments are 84-02.04.2020.A073 and 84-02.04.2018.A234. C57Bl/6J (wildtype, WT) mice were obtained from Janvier Labs (Saint-Berthevin Cedex, France). Mice were housed in standard cages (constant room temperature and humidity, with 12 h light/dark cycles) and had free access to standard pelleted food and tap water.

### REVs and PLEVs isolation

Whole blood (15 ml) from anemic and non-anemic stable coronary arteries disease (CAD) patients was collected in EDTA tubes. To isolate EVs from mice, blood was pooled from three mice yielding a total volume of 3 ml. The blood was centrifuged, and plasma was collected. The buffy coat was removed, and the RBCs were washed three times with phosphate buffered saline (PBS, Sigma-Aldrich, St. Louis, USA). The RBC suspension (40% haematocrit (4 ml) with PBS (6 ml)) was stored at 4°C for 48 h to facilitate the vesiculation. Next, the RBC suspension was centrifuged at 800 x g for 10 min, the supernatant was filtered using a 0.8 *μ*m syringe filter (Corning, New York, USA) to remove large particles such as apoptotic bodies and debris. Next, filtered supernatant was centrifuged at 21,000 x g at 4°C for 45 min and washed with filtered PBS. This centrifugation step was repeated for two times. Plasma vesicles were also isolated following same centrifugation steps.

### Characterization of REVs and PLEVs

#### Dynamic Light Scattering (DLS)

Brownian motion based DLS measurements were used to assess the size distribution of the isolated EVs. As described above, both REVs and PLEVs were isolated from anemic and non-anemic patients, the pellet was dissolved in 1 ml of filtered PBS. The particle size distribution was analysed using the NANOTRAC WAVE II/Q/ZETA (Microtrac, York, USA). Five measurements per sample, each lasting 30 seconds, were performed and then average values were collected using the DIMENSIONS LS software.

#### Nanoparticle tracking analysis (NTA)

NTA was used to determine the size and concentration based on a principle that a laser beam illuminates the particles resulting in scattered light, which is collected by a microscope objective and is viewed with a digital camera. The camera captures a video of the particles moving under Brownian motion. As described above, both REVs and PLEVs were isolated from both anemic and non-anemic patients, diluted in 1000 *μ*l demineralized water, and injected into the chamber of NanoSight NS300 (Malvern Panalytical, Malvern, UK). For each sample, three recordings were made, each lasting 30 seconds. The particle size distribution and concentration were analysed with the Nanosight NTA 1.4 software.

#### Transmission electron microscopy (TEM)

As described above, both PLEVs and REVs were isolated from anemic human and mouse samples. The EVs were fixed by mixing the sample in a 1:4 ratio with fixative (2.5% glutaraldehyde, 4% paraformaldehyde in 0.1M cacodylate buffer, pH 7.4) and incubated for 10 min at RT. Approximately, 5 *μ*l of the mixture was placed on a grid (SP162, pioloform on copper, 200 mesh, Plano, Wetzlar, Germany). Following this, a negative contrast staining with uranyl acetate was performed, where a drop of uranyl acetate was applied and removed three times. Imaging was conducted using the H-7100 TEM (Hitachi, Tokyo, Japan) with a Morada camera (EMSIS, Münster, Germany) and iTEM software (Olympus, Münster, Germany).

### Western Blot

After REVs isolation, the pellet was lysed with 30 *μ*l of radioimmunoprecipitation assay (RIPA) buffer. Afterwards, REVs concentrations of the samples were measured using the DC™ Protein Assay Kit (Bio-Rad, Feldkirchen, Germany) and samples were prepared for WB. The samples were denatured and loaded on a 4%–12% Bis-Tris gradient gel QPAGE™ (Smobio, Hsinchu City, Taiwan). Followed by a transfer onto nitrocellulose membrane and a total protein staining by using Total Protein Stains (LI-COR Biosciences, Bad Homburg, Germany). The total protein staining was detected at 700 nm on the Odyssey® Fc Imaging System (LI-COR Biosciences). Membrane was blocked with intercept TBS Protein-Free Blocking Buffer (LI-COR) for 1 h at RT, followed by overnight incubation of the primary antibody at 4°C. Corresponding secondary antibody was incubated for 1 h at RT and detected at the the Odyssey® Fc Imaging System.

### NO consumption assay

The REV-mediated NO consumption was assessed as previously described [21] with a chemiluminescence detector (CLD 88, Eco Medics, Duernten, Switzerland). Briefly, blood was collected in EDTA tubes and centrifuged at 800 x g for 10 min at 4 °C, the buffy coat was removed, and REVs were isolated from 7 ml of RBC suspension from both groups. The pellet was stored until further use.

First, the system was calibrated by injecting known amounts of nitrite into a reduction solution comprising 45 mmol/L potassium iodide (KI) and 10 mmol/L iodine (I2) in glacial acetic acid, kept in a septum-sealed reaction chamber at 60°C and continuously purged with helium. The reaction chamber was then cleaned with 0.1 M NaOH and ultrapure water. Ten microliters of a freshly prepared stock solution of DetaNONOate (Cayman Chemicals, Ann Arbor, USA; 120 *μ*M) in PBS (pH 7.4) was added to the reaction chamber containing 40 ml of PBS. After the signal for NO release from the NO donor stabilized, REVs resuspended in 50 *μ*L ultrapure water were injected into the reaction chamber. The resulting in a transient decrease in the NO signal indicative of NO consumption. The extent of NO consumption by REVs was calculated by integrating the (negative) areas under the curve of the quenching reaction and comparing them to the (positive) areas for NO generated by the chemical reduction of known amounts of nitrite standards.

### *In vitro* assessment of EVs uptake by endothelial cells

The uptake of EVs by ECs were assessed using the PKH67 green Fluorescent Cell Linker Kit (Sigma-Aldrich) following the company’s instructions. REVs and PLEVs were prepared as described before and labelled. Briefly, the EV pellet was resuspended in 1 ml PBS and the suspension was centrifuged again at 21000 x g for 30 min at 4°C. The supernatant was removed, 0.5 ml of diluent C was added to the EV pellet, and resuspended. A dye solution with 2 *μ*l of PKH67 and 0.5 ml of diluent C was prepared, added to the EV suspension, and incubated for 5 min. The staining was stopped by adding 1 ml of 1 % BSA in PBS solution for 1 min. After that, the EV suspension was centrifuged for 30 min at 21000 x g and the supernatant was removed. The centrifugation step was repeated, and the supernatant was discarded. The pellet was then dissolved in 100 *μ*l serum-free Human umbilical vein endothelial cells (HUVEC) medium and incubated with HUVECs (Lonza, Basel, Switzerland). After 24 h incubation, the cells were fixed with 4% paraformaldehyde (PFA) (Thermo Fisher Scientific, Waltham, USA) and imaged with Leica DM6 M microscope (Leica Microsystems, Wetzlar, Germany).

To verify RBC vesiculation and the uptake of released REVs by endothelial cells, RBC membranes were labeled with the PKH67 Green Fluorescent Cell Linker Kit (Sigma Aldrich) according to the manufacturer’s protocol. Briefly, the RBC pellet was prepared by centrifuging whole blood at 800 x g for 10 minutes at 4°C. The buffy coat was then removed, and the pellet was washed three times with PBS. Then, 4 ml of diluent C was added to the RBC pellet and resuspended. A dye solution containing 16 *μ*l of PKH67 and 4 ml of diluent C was prepared. Dye solution was added to the RBC suspension and incubated for 10 min. The staining was stopped by adding 8 ml of 1% BSA PBS solution for 1 minute. Next, the RBC suspension was centrifuged at 800 x g for 30 minutes. The supernatant was removed, and the centrifugation step was repeated. The RBC pellet was dissolved in 2 ml of PBS and incubated at 4°C for 48 h to facilitate vesiculation. After incubation, REVs were isolated as mentioned before. The REV pellet was then dissolved in 100 *μ*l of serum-free medium and incubated with HUVECs. After 24 h of incubation, the cells were fixed on slide and imaged with Leica DM6 M microscope (Leica Microsystems).

### *In vitro* studies with isolated aortic rings

To assess the effect of anemic REVs and PLEVs on endothelial function, the EVs were co-incubated with wild-type (WT) mouse aortic rings, and vascular function was evaluated. Both REVs or PLEVs were isolated from anemic and non-anemic CAD patients. The pellet (12.5 *μ*g) was dissolved in 100 *μ*l of serum free HUVEC medium. Mouse thoracic aorta was carefully dissected and separated from surrounding perivascular adipose tissue. The aorta was gently perfused with prepared REVs or PLEVs and placed in a petri dish containing 1.9 ml of serum free HUVEC medium. The aorta was incubated for 18 h at 37°C. After the incubation, the aorta was cut into 2 mm segments, mounted on a wire myograph system (Danish Myo Technology, Aarhus, Denmark) and stretched to 9.8 mN. The segments were allowed to equilibrate for 45 min in Krebs-Ringer bicarbonate-buffered salt solution (KRB, 118.5 NaCl, 4.7 KCl, 2.5 CaCl_2_, 1.2 MgSO_4_, 1.2 KH_2_PO_4_, 25.0 NaHCO_3_ and 5.5 glucose in mmol/L).

After normalization, a concentration-response curve (CRC) for phenylephrine (PHE, 0.001-10 μM, Sigma-Aldrich) and acetylcholine (ACH, 0.001-10 μM, Sigma-Aldrich) was generated in presence of the cyclooxygenase inhibitor indomethacin (INDO, 10 *μ*M). Next, to evaluate the contribution of NO to the relaxation response in the arteries the CRC was repeated in the presence of both INDO and Nω-nitro-arginine methyl ester (L-NAME,100 *μ*M, Sigma-Aldrich) a NOS inhibitor. Additionally, the SMC sensitivity to NO was evaluated by performing a CRC in presence of INDO and L-NAME (100 *μ*M), with the NO donor sodium nitroprusside (SNP, 0.01-10 *μ*M, Sigma-Aldrich).

### Mass spectrometry-based proteomics

Both REVs and PLEVs were isolated from anemic and non-anemic CAD patient samples as described before. The samples were lysed with 2% SDS in 300 mM Tris/HCl, (pH 8.0) sonicated to facilitate the complete lysis for EVs. Then the samples were reduced with 20 mM DTT and alkylated with 300 mM IAA. Next, the lysed REVs were digested (SP3 bead digestion), the resulting peptides will be separated using nano-HPLC and analyzed with Orbitrap Fusion Lumos Tribrid Mass Spectrometer (Thermo Fisher Scientific). Data was processed using Proteome Discoverer software (version 2.4.1.15, Thermo Fisher Scientific). RAW files were searched against the human SwissProt database and the MaxQuant contaminant database. A precursor mass tolerance of 10 ppm and a fragment mass tolerance of 0.5 Da were applied. The following modifications were considered: methionine oxidation, N-terminal acetylation, N-terminal methionine loss, and N-terminal methionine loss combined with acetylation as variable modifications, and carbamidomethylation as a static modification. Tryptic cleavage specificity was set with a maximum of two missed cleavage sites.

### Flow cytometry analysis of ROS

To analyse the ROS production in endothelial cell after co-incubation with both REVs and PLEVs, the blood and EVs were prepared as mentioned before. REVs and PLEVs were co-incubated with HUVECs in a 96-well plate for 18 h. After co-incubation, HUVECs were washed once with PBS, trypsinized and stained using the following antibodies and dyes: CD31-V450-PB (clone REA1028, Miltenyi Biotec), ViaKrome 808 Fixable Viability Dye-IR885 (Beckman Coulter, Marseille, France), and 2’,7’-dichlorodihydrofluorescein diacetate (H2DCFDA)-B225-FITC (Thermo Fisher) and analysed using flow cytometer. The data was extracted using FlowJo software v10.5.3 and analysed using Graphpad Prism 10.0.

### Statistical analysis

In *ex vivo* vascular function experiments, all CRCs for contractile stimuli are expressed as absolute values. Relaxing responses were expressed as percentage reduction of the level of contraction. Individual CRCs were fitted to a non-linear sigmoid regression curve (Graphpad Prism 10.0). Sensitivity (pEC_50_), maximal effect (E_max_) are shown as means ± SEM. In all experiments two groups were compared by unpaired t-tests. Two-way ANOVA followed by a Bonferroni post-hoc test was used to compare multiple groups.

## Results

### Characterization of RBC-derived large extracellular vesicles (REVs)

To verify whether REVs play a crucial role in mediating ED, we first established a new method for isolating REVs. One of the major challenges in EV isolation is obtaining a sufficient quantity of EVs from a given volume of blood sample (whether human or mouse) for experimental purposes. To address this, we prepared a suspension of RBCs in PBS and incubated the suspension for 48 h at 4 °C. The REVs released into the supernatant were collected and processed by filtration and centrifugation steps (Fig. 1A). We then characterized both PLEVs and REVs using various methods. Dynamic light scattering (DLS) analysis shows that both REVs and PLEVs ranged in size from 100 to 1000 nm (Fig. 1B), with REVs showing a peak at approximately 200 nm and PLEVs at around 150 nm. The size distribution was consistent between anemic and non-anemic patients, indicating that isolated REVs and PLEVs were categorized as large EVs (Fig. 1B).

**Figure 1:**
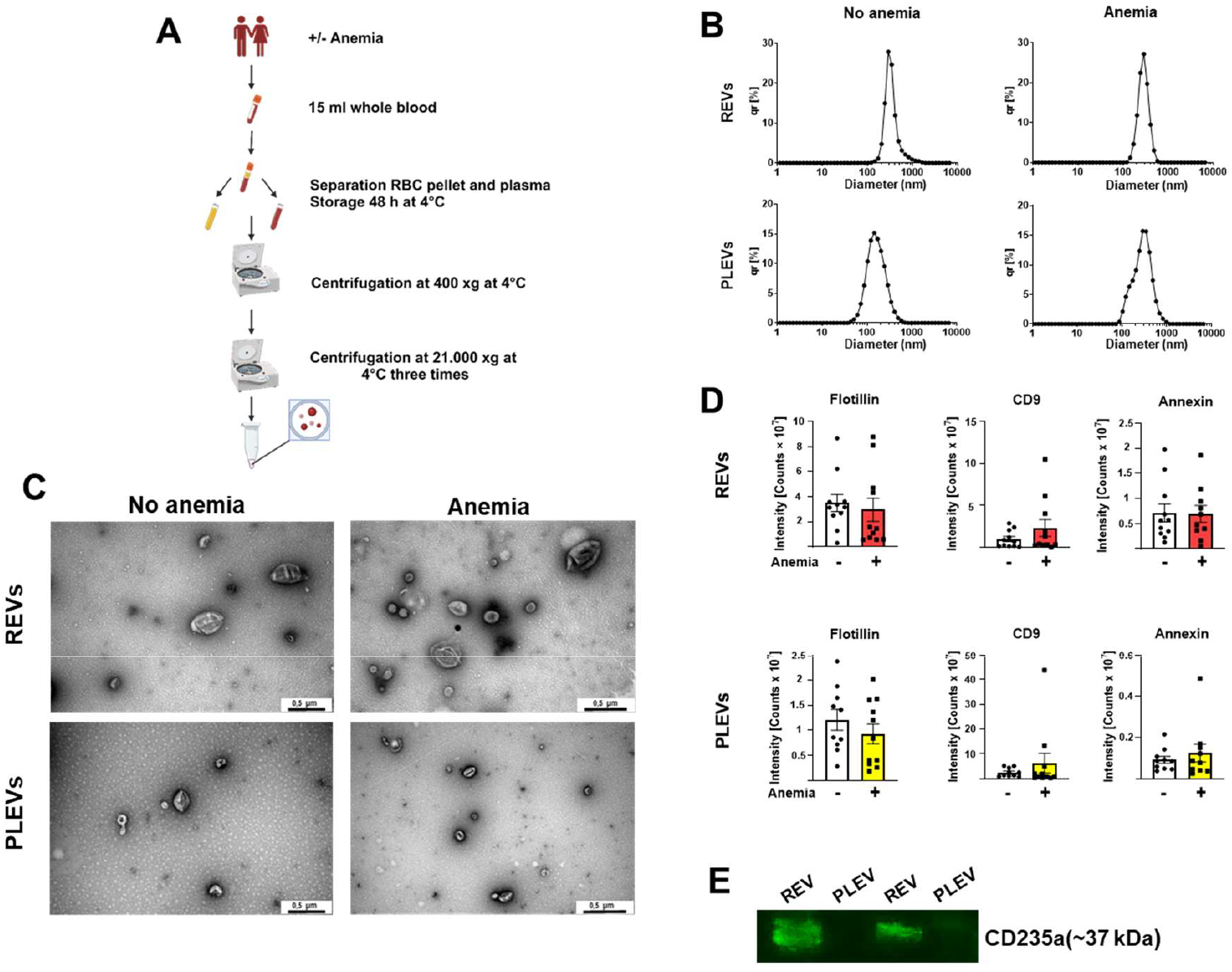
Characterization of RBC-derived (REVs) and plasma derived extracellular vesicles (PLEVs) isolated from anemic and non-anemic CAD patients. **(A)** Isolation protocol for REVs and PLEVs from whole blood. **(B)** Representative size distribution of REVs and PLEVs in non-anemic and anemic CAD patients. **(C)** Representative TEM images of EVs from anemic and non-anemic patients. **(D)** Presence of EV markers: flotillin, CD9, and annexin in PLEVs and REVs isolated from anemic and non-anemic patients. The abundance data is derived from proteomic data. The units ‘‘counts’’ refers to the number of ions detected over a specific period. **(E)** Protein expression of the RBC marker CD235a in lysed REVs and PLEVs.

TEM images further supported these findings, showing a higher number of vesicles per visual field in anemic CAD patients compared to non-anemic controls (Fig. 1C), suggesting that anemia is associated with increased vesicle release. Furthermore, mass spectrometry-based proteome analysis detected EV-specific markers, such as flotillin, CD9, and annexin in both, REVs and PLEVs (Fig. 1D), providing additional layer of evidence that isolated EVs from both sources are pure and contamination-free. Finally, we confirmed the origin of REVs using the RBC-specific marker, glycophorin A (CD235a), which was enriched in REVs but not in PLEVs (Fig. 1E), suggesting a pure EV population in our isolates.

### Increased release of REVs promotes ED by cellular crosstalk

Nanoparticle tracking analysis (NTA) revealed increased vesicle release from RBCs of anemic patients compared to those of non-anemic patients (Fig. 2A), while PLEVs concentrations remained unchanged (Fig. 2B). We also observed increased NO consumption in REVs from anemic patients compared to those from non-anemic individuals suggesting that anemic REVs play a crucial role in regulating NO bioavailability (Fig. 2C). PLEVs only had comparably marginal NO consumption that did not differ between anemic status of the patient.

**Figure 2:**
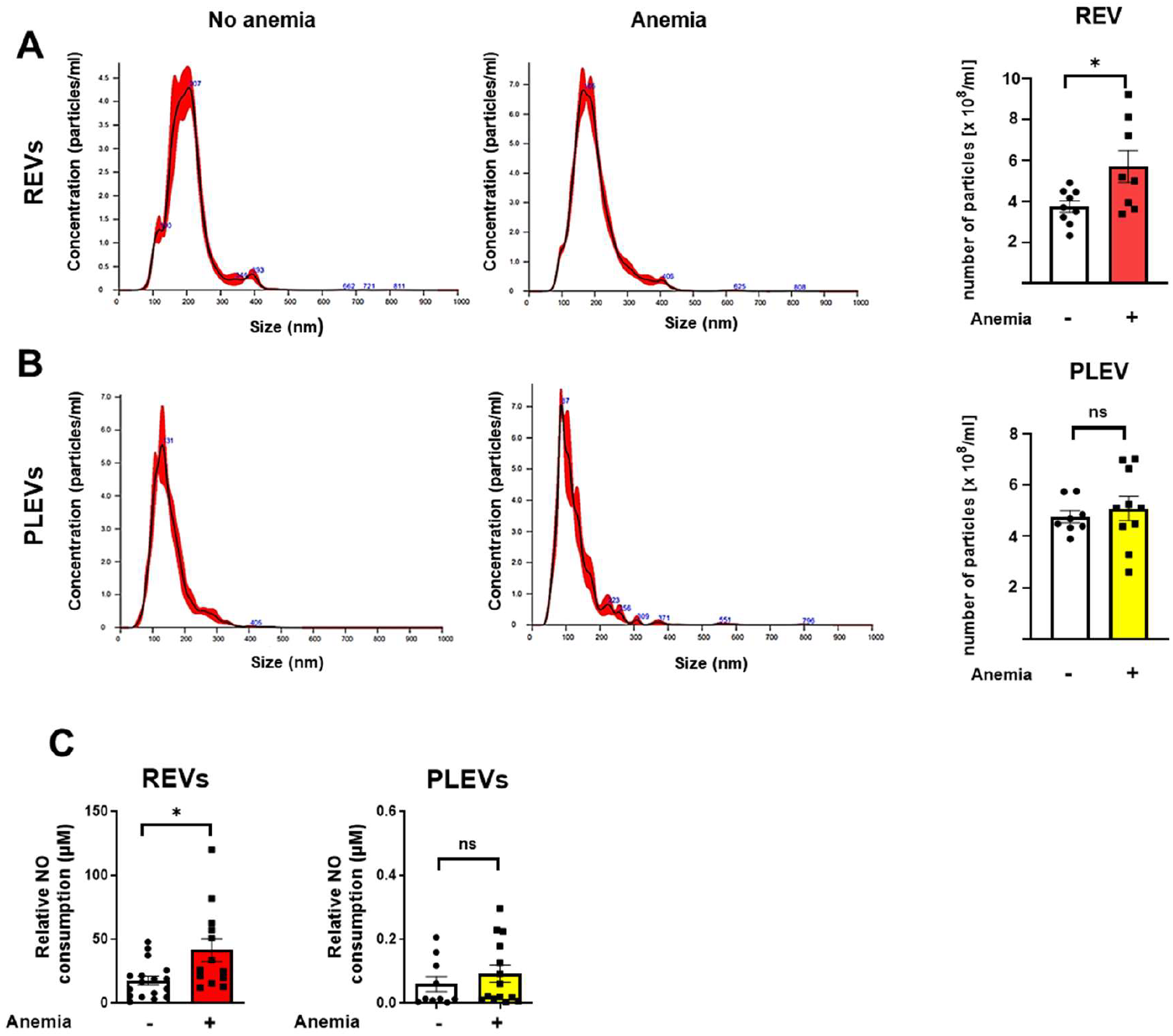
Anemic RBCs show increased vesicle release and enhanced nitric oxide (NO) consumption. REVs (red bar) and PLEVs (yellow bar) were isolated from anemic and non-anemic (white bar) stable coronary artery disease (CAD) patients. Particle number and size distribution was quantified using nanoparticle tracking analysis. **(A-B)** Size distribution and quantification of REVs and PLEVs. **(C)** NO consumption assay in isolated REVs and PLEVs. All values are mean values ± SEM (n = 9-10 per group). *, P ≤ 0.05. Student t-test was used to compare the particles concentration between anemic and non-anemic patients.

Considering that EVs are internalized by recipient cells [11], we examined whether the isolated REVs and PLEVs were taken up by endothelial cells under *in vitro* conditions. After 18 h of co-incubation, PKH67 (a lipophilic fluorescent dye staining the lipid-bilayered membrane of EVs)-labeled REVs and PLEVs were taken up and widely distributed in HUVECs (Fig. 3A-B). Of note, both REVs and PLEVs were also taken up in a dose-dependent manner (Fig. 3C). Additionally, RBCs were stained with PKH67, and vesiculation was allowed for 48 h. Subsequently, the isolated REVs, that contained membrane labelling from the parental cells, were incubated with HUVECs. As expected, after 18 h of co-incubation, PKH67-labeled REVs were widely distributed in HUVECs (Fig. 3D). This further confirms that REVs are taken up by recipient cells, such as EC. In addition, uptake efficacy of both PLEVs and REVs did not depend on whether the EVs were isolated from anemic or non-anemic patients (Suppl. Fig. 1).

**Figure 3:**
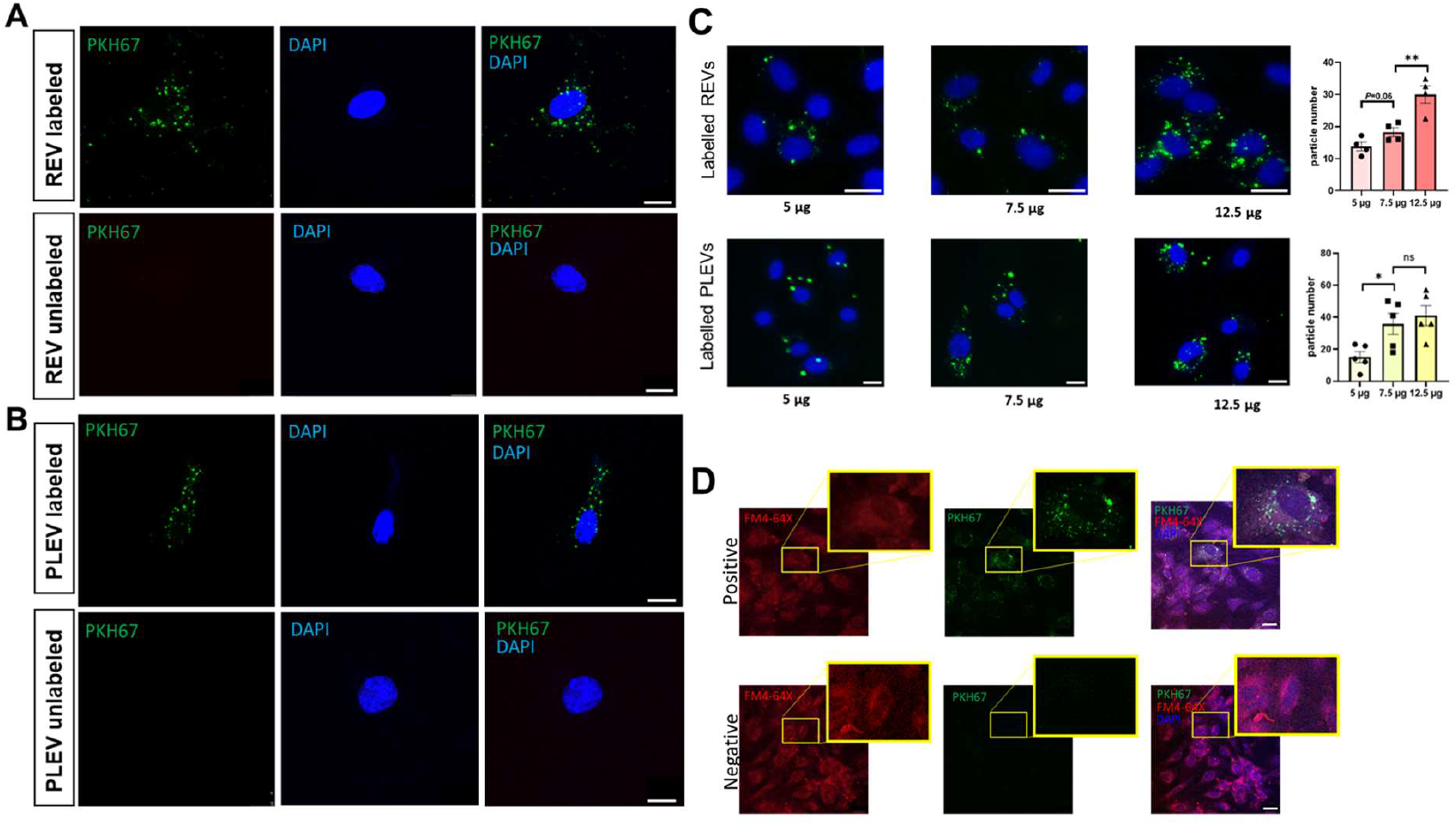
Both RBC-derived (REVs) and plasma-derived extracellular vesicles (PLEVs) are taken up by endothelial cells. Both REVs and PLEVs were isolated from stable coronary artery disease (CAD) patients. **(A-B)** Representative images of HUVECs incubated with either REVs or PLEVs labelled with PKH67 (membrane stain). Nuclei of HUVECs were stained with DAPI. Unlabelled EVs from same patients are used as negative controls. **(C)** Dose-dependent uptake of REVs and PLEVs by HUVECs. Representative images of HUVECs incubated with REVs and PLEVs labelled with PKH67 (membrane stain) of indicated concentrations. Nuclei of HUVECs were stained with DAPI. Scale bar 20 μm. **(D)** Representative images of HUVECs incubated with REVs derived from PKH67-labeled RBCs. EVs from unlabelled RBCs of the same patients served as negative controls. The membrane was stained with FM4-64X, and the nuclei were stained with DAPI. Scale bar is 10 μm. All values are mean values ± SEM. *, P ≤ 0.05. One-way ANOVA was used for multiple condition comparisons.

To evaluate the role of anemic REVs in mediating ED, we assessed both endothelium-dependent and -independent relaxation responses in murine aortic segments that were co-incubated with REVs from anemic and non-anemic CAD patients. In the presence of indomethacin, aortic rings incubated with REVs from anemic patients showed significantly increased contractile responses to phenylephrine (E_max_: 8.72 ± 0.88%) compared to non-anemic controls (E_max_: 6.07 ± 0.61%) (Suppl. Table 1). Additionally, endothelium-dependent relaxation responses to acetylcholine, in the presence of indomethacin, were significantly reduced in aortic segments treated with anemic REVs (E_max_: 48.90 ± 13.18%) compared to non-anemic REVs (E_max_: 81.09 ± 5.14%), indicating impaired endothelial-dependent relaxation (Fig. 4A, Suppl. Table 1). The relaxation responses were equally and completely inhibited in the presence of the NOS inhibitor L-NAME (100 *μ*M), confirming that these relaxation responses are entirely mediated by NO (Fig. 4B, Suppl. Table 1). The relaxation responses to SNP (NO donor) were similar in both groups (Fig. 4C, Suppl. Table 1).

**Figure 4:**
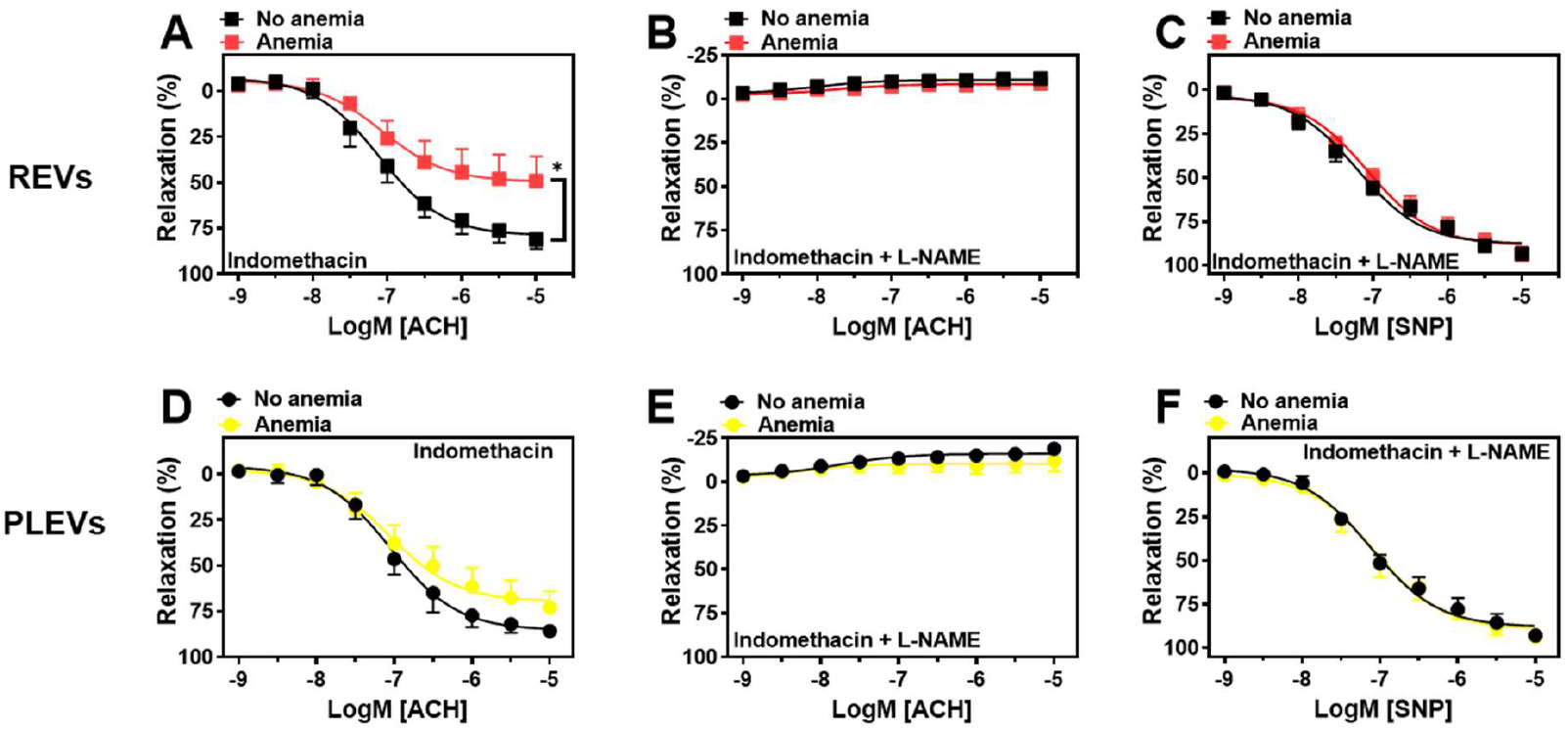
RBC-derived extracellular vesicles (REVs) from anemic CAD patients induce endothelial dysfunction in murine aortic segments. REVs **(A-C**) and plasma-derived extracellular vesicles (PLEVs) **(D-F)** were isolated from anemic and non-anemic stable coronary artery disease (CAD) patients and incubated with murine aortic rings. After 18 h, segments were mounted in a wire myograph system. **(A, D)** Relaxation responses to acetylcholine (ACH) in the presence of indomethacin. **(B, E)** Relaxation responses to ACH in the presence of indomethacin and L-NAME. **(C, F)** Relaxation responses to sodium nitroprusside (SNP, 10 nM-10 μM) in the presence of indomethacin and L-NAME. Values are shown as means ± SEM (n = 6-7 per group). *, P ≤ 0.05. CRCs were analyzed by Two-Way ANOVA and Bonferroni’s post-hoc test. CAD: stable coronary artery disease.

Furthermore, we investigated the effects of PLEVs from anemic and non-anemic patients on endothelial function. Similar to the REVs experiment, PLEVs from both anemic and non-anemic patients were co-incubated with mouse aortic segments for 18 h. The contractile responses of the aortic rings incubated with PLEVs were comparable between the anemic and non-anemic groups (Suppl. Table 1). Likewise, the relaxation responses in the presence of indomethacin were similar for both aortic segments incubated with PLEVs from anemic and non-anemic CAD patients (Fig. 4D, Suppl. Table 1). In the presence of L-NAME, the relaxation responses were equally and completely inhibited in both groups (Fig. 4E, Suppl. Table 1). Additionally, the relaxation responses to SNP were similar in both groups (Fig. 4F, Suppl. Table 1). In summary, these results indicate that REVs, but not PLEVs, from anemic CAD patients contribute to ED.

### REVs and PLEVs from anemic patients carry a diverse array of proteins, including those associated with redox signalling

To gain deeper insights into the content of REVs and PLEVs, we performed an unbiased proteomic analysis. Due to the partly uneven distribution of proteins in isolated EV fractions, we applied specific filter criteria for the analysis. Only proteins that have non-zero abundance in at least five out of eleven patients in one of the groups were considered for further analysis. After filtration, 2369 out of 2389 total proteins detected in REVs were included in the analysis. In the case of PLEVs, 2044 out of 2071 total proteins were considered after filtration.

The volcano plots represent all proteins, including differentially regulated proteins in REVs and PLEVs (Fig. 5A). Next, we selected all differentially expressed proteins in REVs and PLEVs. When comparing anemic to non-anemic patients, approximately 30 proteins were differentially expressed in REVs (Fig. 5B) and 60 proteins in PLEVs (Fig. 5C), revealing differences in the EV cargo content between anemic and non-anemic patient groups. Among these differentially regulated proteins, in REVs, proteins such as heparanase (HPSE), coproporphyrinogen oxidase (CPOX), ATP synthase inhibitory factor subunit 1 (ATP5IF1), and serine hydroxymethyltransferase 2 (SHMT2) are consistently abundant in the anemic REVs of the majority of patients compared to non-anemic CAD patients. Similarly, in PLEVs, talin 2 (TLN2), transferrin receptor (TFRC), and torsin family 4 member A (TOR4A) are more abundant in the PLEVs of anemic CAD patients, whereas fibroblast growth factor receptor 2 (FGFR2), apolipoprotein A2 (APOA2), ST8 Alpha-N-Acetyl-Neuraminide Alpha-2,8-Sialyltransferase 4 (ST8SIA4), ficolin 3 (FCN3), Family With Sequence Similarity 234 Member A (FAM234A), major histocompatibility complex class II DR Beta 5 (HLA-DRB5), Immunoglobulin Lambda Variable 3-21 (IGLV3), and fibrinogen alpha chain (FGA) are consistently abundant in the PLEVs of the majority of non-anemic patients compared to anemic CAD patients.

**Figure 5:**
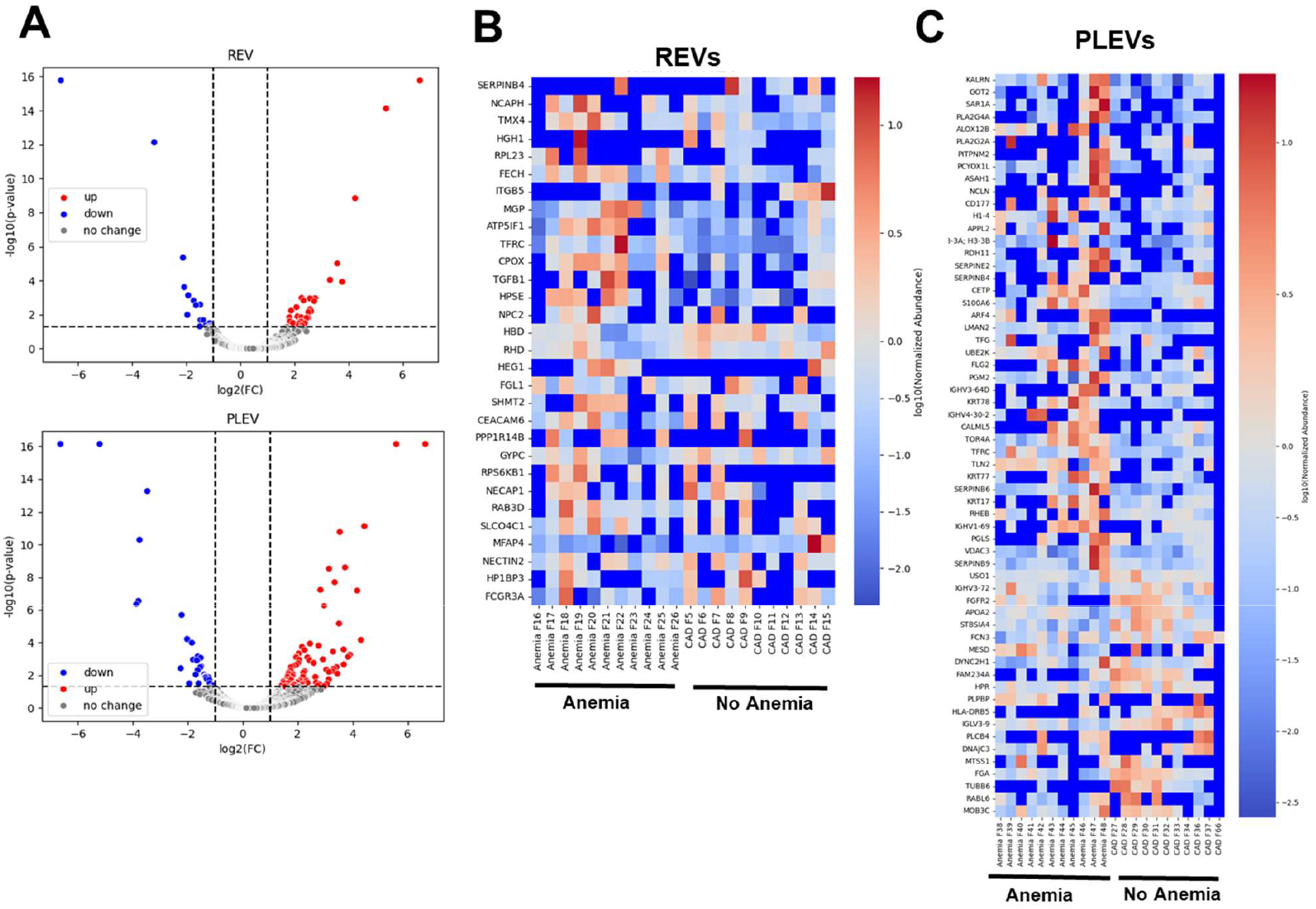
Proteomic analysis of RBC-derived (REVs) and plasma-derived extracellular vesicles (PLEVs) isolated from anemic and non-anemic CAD patients. **(A)** Volcano plots displaying the enrichment of proteins between anemic and non-anemic patients; proteins with occurrence ≥5 in anemic samples or ≥5 in non-anemic samples are considered and selected for further analyses. **(B, C)** Heatmap of the normalized abundances of selected proteins isolated from anemic and non-anemic patients, in REVs **(B)** and PLEVs **(C)**; proteins in the heatmap are ordered according to their enrichment between anemic and non-anemic samples. Non detected proteins were calculated as 0.

Interestingly, anemic REVs showed the specific abundance of proteins such as tubulin beta-2A chain, NADH-cytochrome b5 reductase 1, immunoglobulin kappa variable 1-17, ornithine aminotransferase, and cytochrome b-c1 complex subunit 6 (Suppl. Table 2), which were not detected in non-anemic CAD patients. Similarly, in PLEVs, we observed exportin-7, tetraspanin-15, AP-3 complex subunit beta-1, heat shock protein 75 kDa, ubiquitin carboxyl-terminal hydrolase, and 2-hydroxyacyl-CoA lyase 2, among others, in anemic patients, highlighting their potential role as biomarkers in anemia (Suppl. Table 2).

Since REVs are known to carry redox proteins, we specifically analysed the redox proteins expressed in both REVs and PLEVs and compared them between anemic and non-anemic patients (Fig. 6A-B). This analysis suggests that REVs from anemic patients are associated with an increased average abundance of redox-regulating proteins compared to REVs from non-anemic patients. Due to high variability between samples, these redox proteins did not significantly differ between the anemic and non-anemic patients. Yet, several antioxidants such as superoxide dismutase 1 (SOD1), glutathione peroxidase 1 (GPX1), catalase (CAT), were reduced in anemic REVs compared to those from non-anemic patients (Fig. 6A). Noteworthy, we also observed an increased average abundance of myeloperoxidase (MPO) in anemic REVs compared to non-anemic REVs. No specific pattern in the distribution of redox proteins was observed in PLEVs, except for glutathione S-transferase mu 2 (GSTM2), which was notably enriched in anemic PLEVs compared to non-anemic ones (Fig. 6B). Interestingly, REVs, but not PLEVs, incubated with HUVECs aggravated oxidative stress in HUVECs (Fig. 6C, Suppl. Figure 3). These findings suggest a potential role of REVs redox proteins in promoting endothelial dysfunction (ED) in anemia

**Figure 6:**
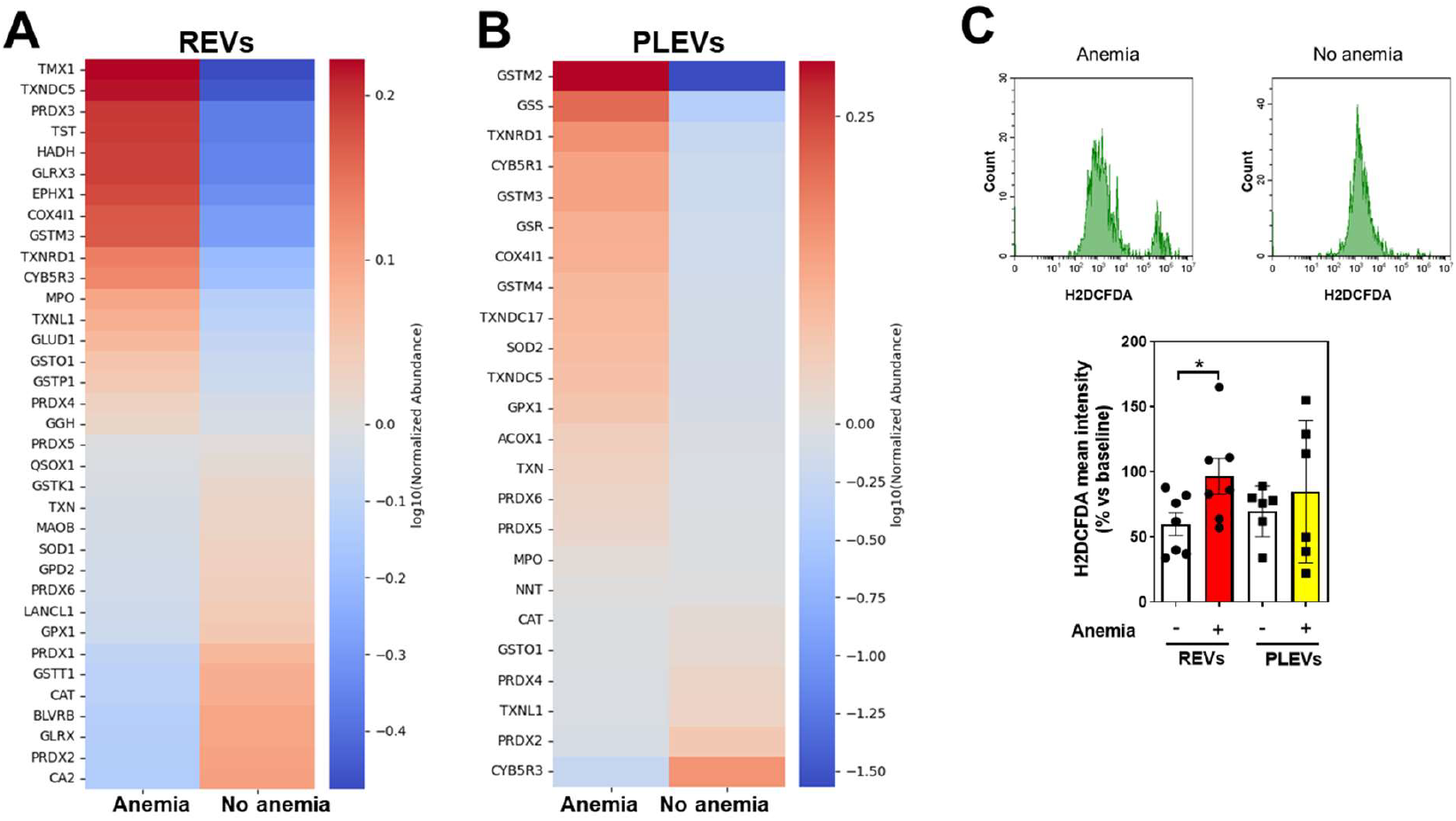
Abundance of redox proteins in RBC-derived (REVs) and plasma-derived extracellular vesicles (PLEVs) isolated from anemic and non-anemic patients. **(A-B**) Heatmap of normalized abundance of selected redox proteins isolated from anemic and non-anemic patients in REVs **(A)** and PLEVs **(B)**. Proteins in the heatmap are ordered according to their enrichment between anemic and non-anemic samples, and are normalized accordance with the average of a protein across different samples is 1. Dark red represents higher abundance, grey indicates no difference, dark blue demonstrates lower abundance. **(C)** Flow cytometry analysis of H2DFCA mean fluorescence in the HUVECS treated with REVs and PLEVs from anemic and non-anemic patients. All values are mean values ± SEM. *, P ≤ 0.05. Student t-test was used to compare between anemic and non-anemic patients.

## Discussion

The field of EV-mediated intercellular crosstalk has attracted significant attention in recent years, particularly regarding its potential applications in non-invasive diagnostics and therapeutic interventions. EVs, which include small EVs (known previously as exosomes), and large EVs (known previously microvesicles), serve as carriers of molecular cargoes, facilitating communication between cells and providing insights into disease states. Their role as biomarkers in various conditions, including cancer and CVD, underscores their relevance in current medical research. However, the specific contributions of EVs in anaemia remain largely unexplored, highlighting an area deserving further investigation. Our study aimed to elucidate the differential effects of REVs and PLEVs from anemic patients on endothelial function, a critical factor in cardiovascular health.

This investigation yielded several key findings that enhance our understanding of the interplay between EVs and endothelial responses. First, we successfully characterized the isolated particles from RBCs and plasma as EVs, confirming their identity and relevance in our study. Notably, we observed an increase in RBC vesiculation among anemic CAD patients compared to non-anemic controls. This finding suggests a potential link between anemia and altered EV production that could serve as a mechanism for impaired endothelial function. Further examination revealed that anemic REVs exhibited increased NO consumption relative to REVs derived from non-anemic CAD patients, indicating a possible enhancement of ED attributed to the altered EV cargo in anemic conditions. Importantly, both PLEVs and REVs were readily taken up by ECs *in vitro*, demonstrating the capacity of these vesicles to influence endothelial behaviour. However, while REVs from anemic CAD patients were shown to induce ED in isolated murine aortic rings following co-incubation, PLEVs did not exhibit this effect, suggesting a distinct functional divergence between EV types in the context of endothelial health. Proteomic analysis further revealed altered abundance of redox proteins in REVs from anemic CAD patients compared to non-anemic individuals. This may provide insights into the oxidative stress mechanisms underpinning ED and highlight the potential of REVs as a focus for future therapeutic strategies.

We established a modified protocol for the isolation of REVs from human samples, incorporating a 48 h storage period of RBC suspension at a controlled cold temperature of 4°C. This incubation step not only fosters RBC maturation but also significantly enhances the yield of REVs compared to freshly isolated RBCs. Previous research has established that exosomes are predominantly released during the maturation of reticulocytes into erythrocytes, whereas microvesicles are primarily secreted by mature RBCs, resulting in a uniform EV profile in the latter [20]. Consequently, our focus was directed towards large EVs rather than exosomes. Notably, NTA analysis indicated a marked increase in vesiculation among RBCs derived from anemic patients, aligning with existing literature that correlates elevated RBC vesiculation with various pathological conditions [20, 22], including cardiovascular and metabolic diseases. However, we did not observe a corresponding increase in PLEVs in the anemic cohort, likely attributable to the diverse origins of EVs present in plasma and the influence of underlying comorbidities on different cell types.

Recent studies have elucidated the role of REVs in the pathogenesis of ED under pathological conditions such as diabetes and intermittent hypoxia [23]. To further investigate the implications of anemic REVs in the context of anemia, we examined their interaction with human endothelial cells, specifically HUVEC cells, through *in vitro* uptake assays. Our results indicate that both REVs and PLEVs are readily internalized by these cells, suggesting the potential for *in vivo* uptake. However, an *in vivo* bio-distribution study is warranted to substantiate these *in vitro* observations. Additionally, co-incubation of REVs with murine aortic rings revealed that exposure to anemic REVs resulted in significantly diminished NO-dependent relaxation responses, in contrast to PLEVs. These findings reinforce the hypothesis that EVs contribute to ED and vascular inflammation by modulating NO production and oxidative stress [23], thereby implicating dysfunctional RBCs as a potential mechanism underlying ED in anemic individuals.

Further studies have demonstrated that large EVs, specifically microparticles or microvesicles derived from RBCs, exhibit significant NO scavenging capabilities, comparable to those of free haemoglobin [24]. Notably, the rate of NO scavenging by these large EVs (microvesicles) is approximately 1000-times greater than that of intact RBCs. This phenomenon raises concerns regarding the implications of RBC hemolysis and microparticle formation, processes that are associated with adverse outcomes in patients receiving older stored blood, primarily due to enhanced NO scavenging [25]. Furthermore, our research indicates that REVs derived from anemic patients show an increased NO consumption. We posit that the augmented vesiculation and associated enhancement in NO consumption may play a critical role in modifying NO production and bioavailability, particularly in the context of anemia *in vivo*.

In our study, we identified a distinct array of proteins that were particularly enriched in anemic patients, indicating their potential utility as biomarkers for anemia-related vascular dysfunction. Notably, among the enriched proteins in anemic REVs were NADH-cytochrome b5 reductase 1, which is integral to the reduction of methemoglobin, and immunoglobulin kappa variable 1-17, a key player in the immune response [26, 27]. Additionally, in PLEVs, we observed an enrichment of AP-3 complex subunit beta-1 and tetraspanin-15, both of which are implicated in vesicle formation, alongside other significant proteins such as heat shock protein 75 kDa and ubiquitin carboxyl-terminal hydrolase. However, the precise roles of these proteins in recipient cells, as well as the underlying mechanisms for their selective enrichment in anemic patients, remain to be elucidated and warrant further comprehensive investigation.

As demonstrated in previous studies, promoting oxidative stress may serve as a potential mechanism by which RBCs contribute to ED in anemia [18]. Our investigation aligns with previous research on REVs [23], revealing an increased abundance of proteins associated with redox homeostasis, including glutathione S-transferase, thioredoxin, and peroxiredoxin in REV cargo from anemic subjects. Notably, the presence of myeloperoxidase (MPO), an enzyme linked to vascular endothelial damage and the enhancement of reactive oxygen species (ROS) formation, further underscores the potential for oxidative stress to diminish NO bioavailability in this context [28]. Additionally, reductions in key antioxidant enzymes, such as superoxide dismutase (SOD), glutathione peroxidase 1 (GPX1), and catalase (CAT) in anemic REVs compared to their non-anemic counterparts, suggest a potential redox imbalance that warrants further investigation. These findings not only reveal the altered redox regulation in RBCs from anemic patients but also highlight crucial targets for future research aimed at understanding the specific mechanisms underlying these phenomena.

Our data indicate a significant association between anemia and heightened vesiculation of REVs in patients with CAD, in contrast to PLEVs. The REVs derived from anemic subjects exhibit a marked increase in NO consumption, potentially influencing systemic NO bioavailability. Co-incubation experiments further elucidate that only anemic REVs, and not PLEVs, are capable of inducing ED in isolated murine aortic rings. Additionally, REVs obtained from anemic patients demonstrate an elevated presence of various redox proteins, including oxidative stress-promoting enzymes such as MPO. Additionally, anemic REVs but not PLEVs induce enhanced ROS production in ECs. Collectively, these findings underscore the role of anemic REVs in contributing to ED through NO consumption and modulation of the endothelial redox state.

## Data availability statement

All the raw data supporting results and conclusions of this manuscript will be made available by the corresponding author, without undue reservation.

## Acknowledgments

The authors wish to thank Stefanie Becher for technical assistance with animal experiments and Susanne Pfeiler for organisational help.

## Author Contributions

RC designed the study. IS, VY, LH, ST, PK, AB, and MRH contributed to EV isolation and characterization. IS and RC contributed to the myograph experiments. PW arranged the CAD patient samples. RC and IS contributed to immunohistochemistry. AS conducted the mass spectrometry experiments. TP, RC, AL, and MRH contributed to proteomics analysis. RC and MRH, IS interpreted the data and drafted the manuscript. MK, NG, CJ, and RC secured funding. GN, MRH, CJ, NG, MK, and RC critically revised the manuscript. All authors contributed to the article and approved the final version for submission.

## References

1. Alaarg, A., et al., Red blood cell vesiculation in hereditary hemolytic anemia. Front Physiol, 2013. 4: p. 365.

2. Erdbrugger, U. and J. Lannigan, Analytical challenges of extracellular vesicle detection: A comparison of different techniques. Cytometry A, 2016. 89(2): p. 123–34.

3. Konoshenko, M.Y., et al., Isolation of Extracellular Vesicles: General Methodologies and Latest Trends. Biomed Res Int, 2018. 2018: p. 8545347.

4. Villysson, A., A. Tontanahal, and D. Karpman, Microvesicle Involvement in Shiga Toxin-Associated Infection. Toxins (Basel), 2017. 9(11).

5. Thery, C., M. Ostrowski, and E. Segura, Membrane vesicles as conveyors of immune responses. Nat Rev Immunol, 2009. 9(8): p. 581–93.

6. Chiangjong, W., et al., Red Blood Cell Extracellular Vesicle-Based Drug Delivery: Challenges and Opportunities. Front Med (Lausanne), 2021. 8: p. 761362.

7. Kumar, M.A., et al., Extracellular vesicles as tools and targets in therapy for diseases. Signal Transduct Target Ther, 2024. 9(1): p. 27.

8. Arroyo, J.D., et al., Argonaute2 complexes carry a population of circulating microRNAs independent of vesicles in human plasma. Proc Natl Acad Sci U S A, 2011. 108(12): p. 5003–8.

9. Tabet, F., et al., HDL-transferred microRNA-223 regulates ICAM-1 expression in endothelial cells. Nat Commun, 2014. 5: p. 3292.

10. Tkach, M. and C. Thery, Communication by Extracellular Vesicles: Where We Are and Where We Need to Go. Cell, 2016. 164(6): p. 1226–1232.

11. Kwok, Z.H., C. Wang, and Y. Jin, Extracellular Vesicle Transportation and Uptake by Recipient Cells: A Critical Process to Regulate Human Diseases. Processes (Basel), 2021. 9(2).

12. Jansen, F., G. Nickenig, and N. Werner, Extracellular Vesicles in Cardiovascular Disease: Potential Applications in Diagnosis, Prognosis, and Epidemiology. Circ Res, 2017. 120(10): p. 1649–1657.

13. Hosen, M.R., et al., Circulating MicroRNA-122-5p Is Associated With a Lack of Improvement in Left Ventricular Function After Transcatheter Aortic Valve Replacement and Regulates Viability of Cardiomyocytes Through Extracellular Vesicles. Circulation, 2022. 146(24): p. 1836–1854.

14. Zhang, X., et al., Extracellular Vesicles in Cardiovascular Diseases: Diagnosis and Therapy. Front Cell Dev Biol, 2022. 10: p. 875376.

15. Pernow, J., et al., Red blood cell dysfunction: a new player in cardiovascular disease. Cardiovasc Res, 2019. 115(11): p. 1596–1605.

16. Kuhn, V., et al., Red Blood Cell Function and Dysfunction: Redox Regulation, Nitric Oxide Metabolism, Anemia. Antioxid Redox Signal, 2017. 26(13): p. 718–742.

17. Wischmann, P., et al., Anaemia is associated with severe RBC dysfunction and a reduced circulating NO pool: vascular and cardiac eNOS are crucial for the adaptation to anaemia. Basic Res Cardiol, 2020. 115(4): p. 43.

18. Chennupati, R., et al., Chronic anemia is associated with systemic endothelial dysfunction. Front Cardiovasc Med, 2023. 10: p. 1099069.

19. Ma, S.R., et al., Red Blood Cell-Derived Extracellular Vesicles: An Overview of Current Research Progress, Challenges, and Opportunities. Biomedicines, 2023. 11(10).

20. Thangaraju, K., et al., Extracellular Vesicles from Red Blood Cells and Their Evolving Roles in Health, Coagulopathy and Therapy. Int J Mol Sci, 2020. 22(1).

21. Quast, C., et al., Aortic Valve Stenosis Causes Accumulation of Extracellular Hemoglobin and Systemic Endothelial Dysfunction. Circulation, 2024. 150(12): p. 952–965.

22. Yang, L., et al., Roles and Applications of Red Blood Cell-Derived Extracellular Vesicles in Health and Diseases. Int J Mol Sci, 2022. 23(11).

23. Zhang, L., et al., Extracellular vesicles and endothelial dysfunction in infectious diseases. J Extracell Biol, 2024. 3(4): p. e148.

24. Jomova, K., et al., Several lines of antioxidant defense against oxidative stress: antioxidant enzymes, nanomaterials with multiple enzyme-mimicking activities, and low-molecular-weight antioxidants. Arch Toxicol, 2024. 98(5): p. 1323–1367.

25. Liu, C., et al., Nitric oxide scavenging by red cell microparticles. Free Radic Biol Med, 2013. 65: p. 1164–1173.

26. Saleh, M.C. and S. McConkey, NADH-dependent cytochrome b5 reductase and NADPH methemoglobin reductase activity in the erythrocytes of Oncorhynchus mykiss. Fish Physiol Biochem, 2012. 38(6): p. 1807–1813.

27. Engelbrecht, E., et al., Resolving haplotype variation and complex genetic architecture in the human immunoglobulin kappa chain locus in individuals of diverse ancestry. Genes Immun, 2024. 25(4): p. 297–306.

28. Poisson, J., et al., Erythrocyte-derived microvesicles induce arterial spasms in JAK2V617F myeloproliferative neoplasm. J Clin Invest, 2020. 130(5): p. 2630–2643.

