## Supplementary Figures and tables for "Large extracellular vesicles derived from red blood cells in coronary artery disease patients with anemia promote endothelial dysfunction"

### Slide 1
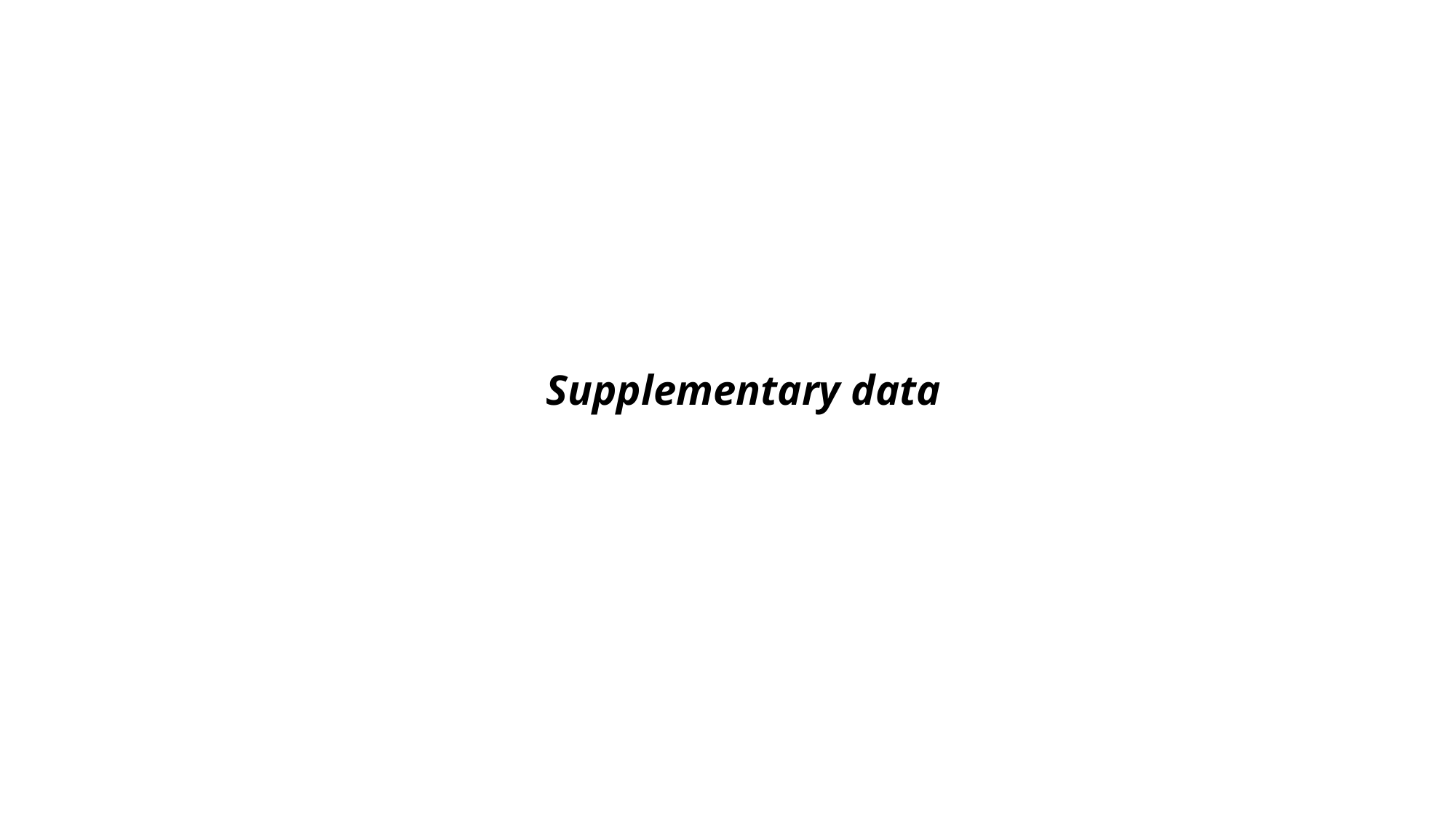

#### Supplementary data

### Slide 2
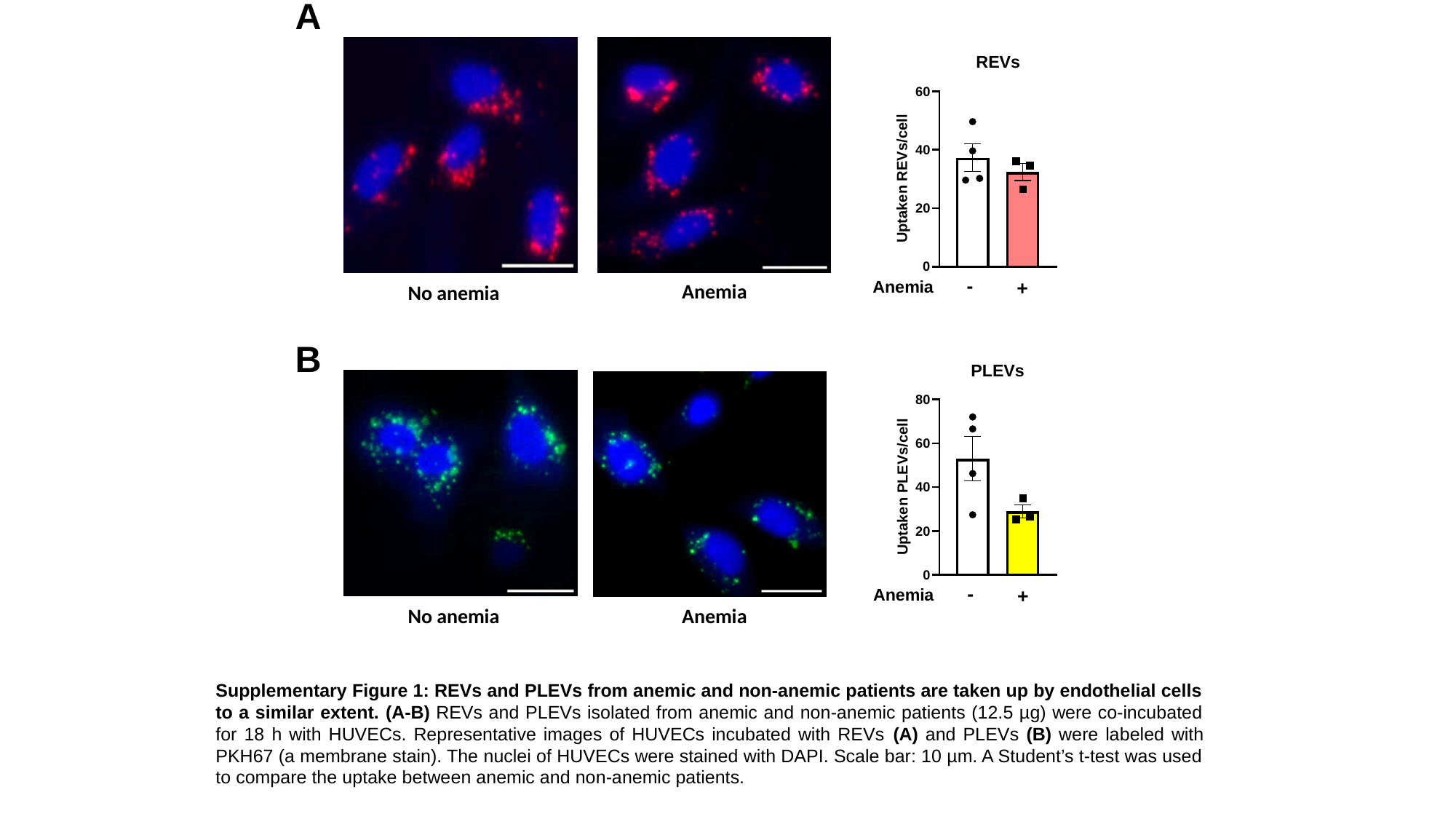

A
Anemia
No anemia
B
No anemia
Anemia
Supplementary Figure 1: REVs and PLEVs from anemic and non-anemic patients are taken up by endothelial cells to a similar extent. (A-B) REVs and PLEVs isolated from anemic and non-anemic patients (12.5 µg) were co-incubated for 18 h with HUVECs. Representative images of HUVECs incubated with REVs (A) and PLEVs (B) were labeled with PKH67 (a membrane stain). The nuclei of HUVECs were stained with DAPI. Scale bar: 10 µm. A Student’s t-test was used to compare the uptake between anemic and non-anemic patients.

### Slide 3
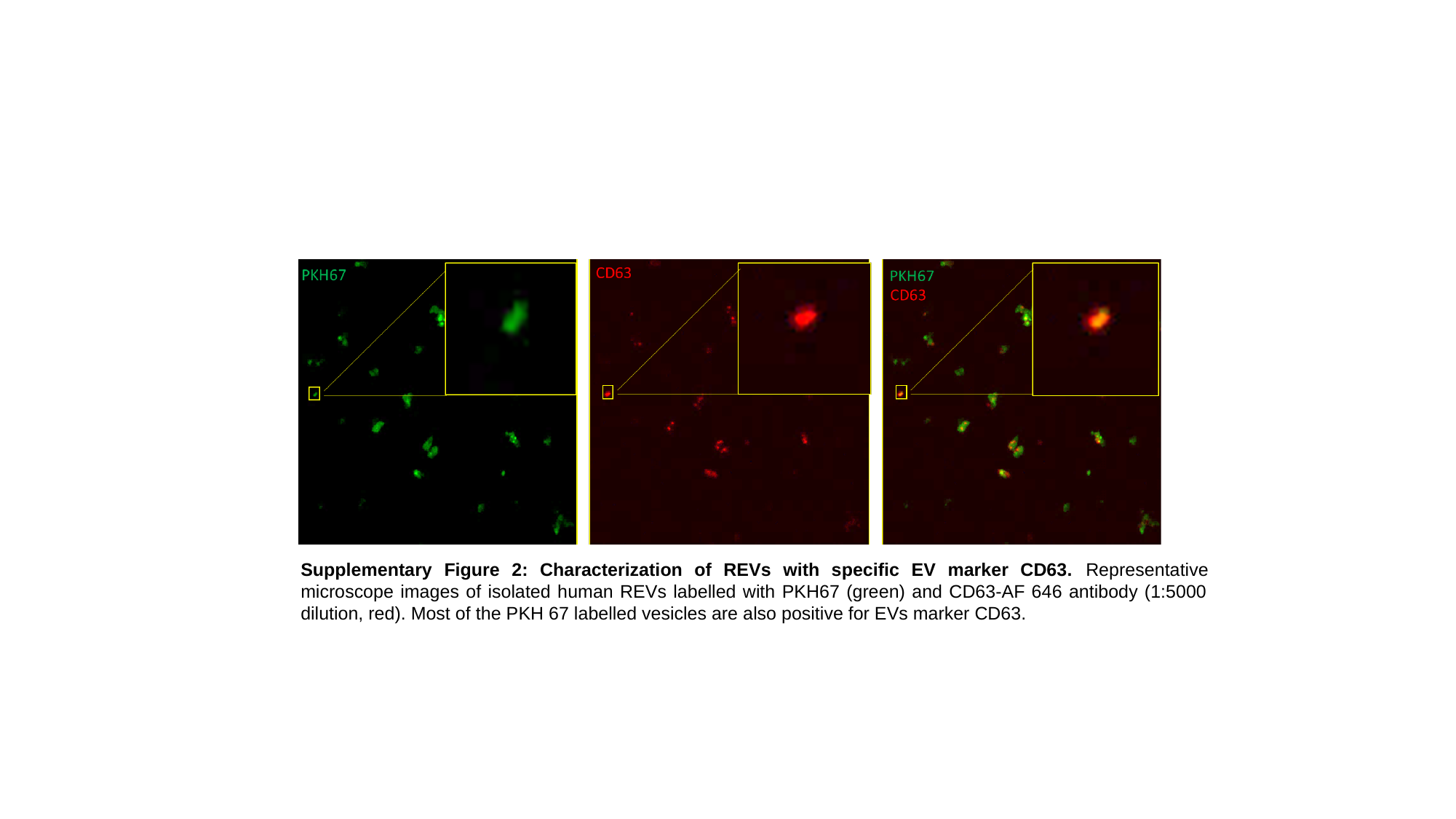

Supplementary Figure 2: Characterization of REVs with specific EV marker CD63. Representative microscope images of isolated human REVs labelled with PKH67 (green) and CD63-AF 646 antibody (1:5000 dilution, red). Most of the PKH 67 labelled vesicles are also positive for EVs marker CD63.

### Slide 4
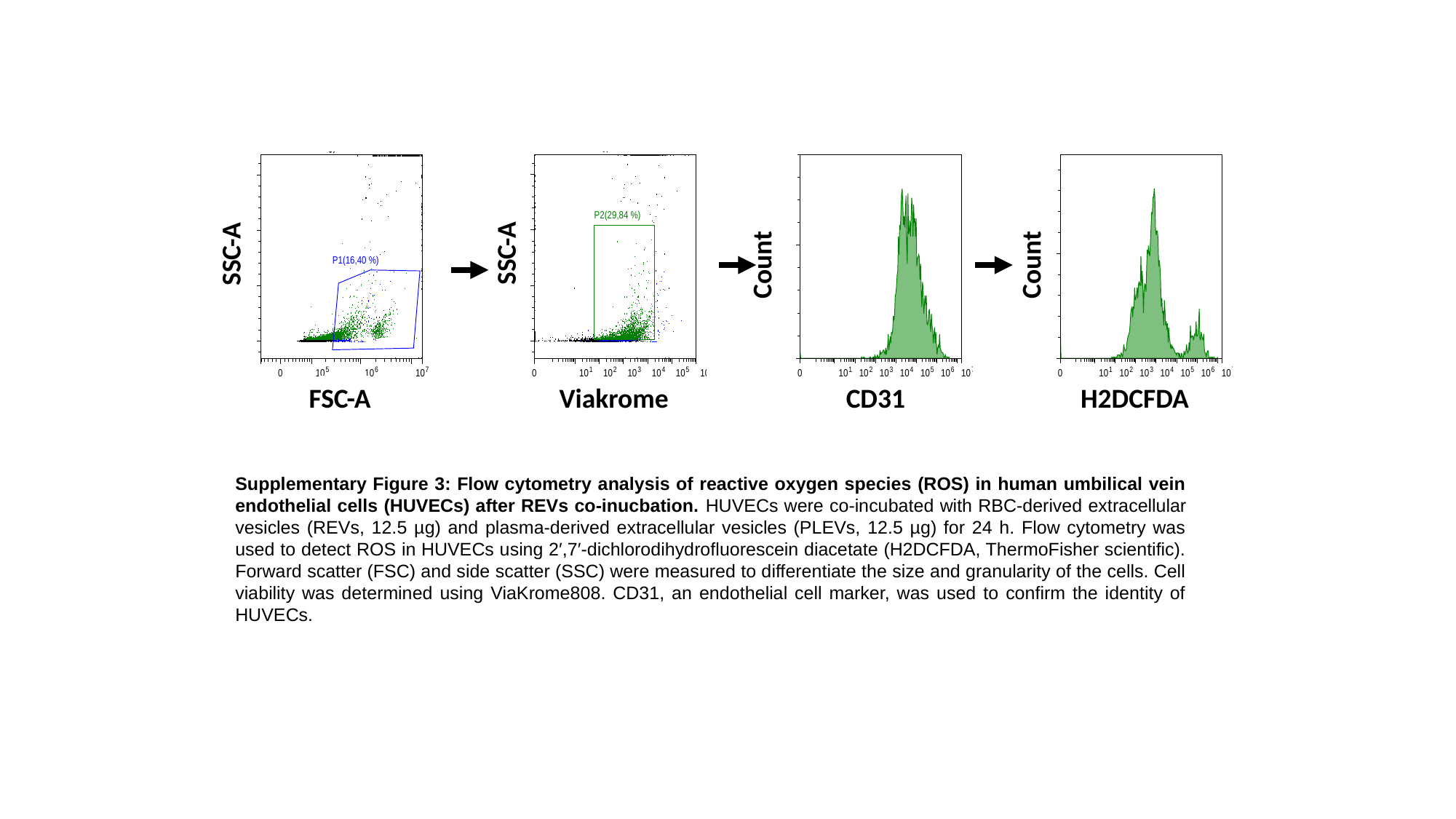

SSC-A
SSC-A
Count
Count
Viakrome
CD31
H2DCFDA
FSC-A
Supplementary Figure 3: Flow cytometry analysis of reactive oxygen species (ROS) in human umbilical vein endothelial cells (HUVECs) after REVs co-inucbation. HUVECs were co-incubated with RBC-derived extracellular vesicles (REVs, 12.5 µg) and plasma-derived extracellular vesicles (PLEVs, 12.5 µg) for 24 h. Flow cytometry was used to detect ROS in HUVECs using 2′,7′-dichlorodihydrofluorescein diacetate (H2DCFDA, ThermoFisher scientific). Forward scatter (FSC) and side scatter (SSC) were measured to differentiate the size and granularity of the cells. Cell viability was determined using ViaKrome808. CD31, an endothelial cell marker, was used to confirm the identity of HUVECs.

### Slide 5
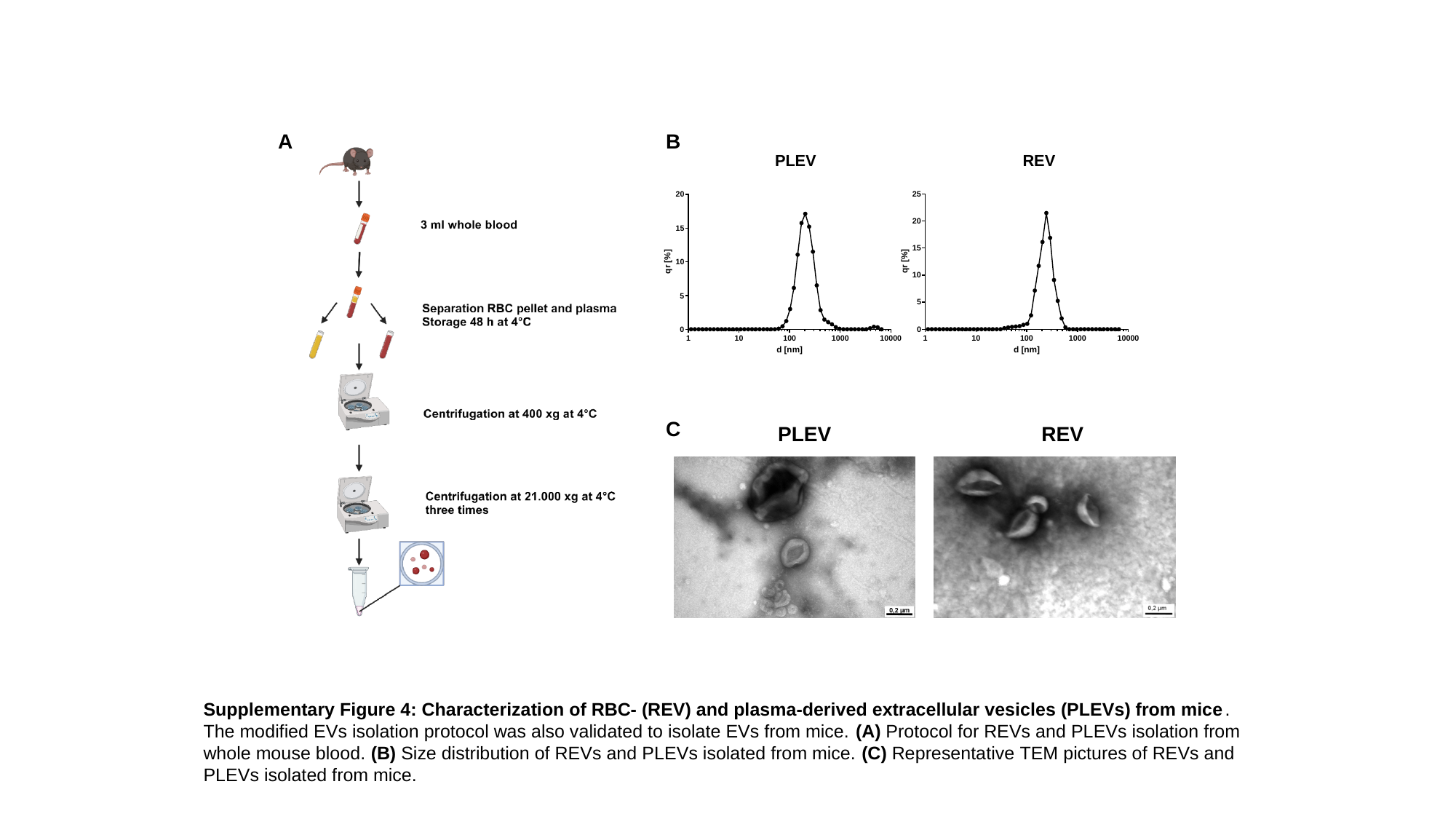

A
B
REV
PLEV
C
PLEV
REV
Supplementary Figure 4: Characterization of RBC- (REV) and plasma-derived extracellular vesicles (PLEVs) from mice. The modified EVs isolation protocol was also validated to isolate EVs from mice. (A) Protocol for REVs and PLEVs isolation from whole mouse blood. (B) Size distribution of REVs and PLEVs isolated from mice. (C) Representative TEM pictures of REVs and PLEVs isolated from mice.

### Slide 6
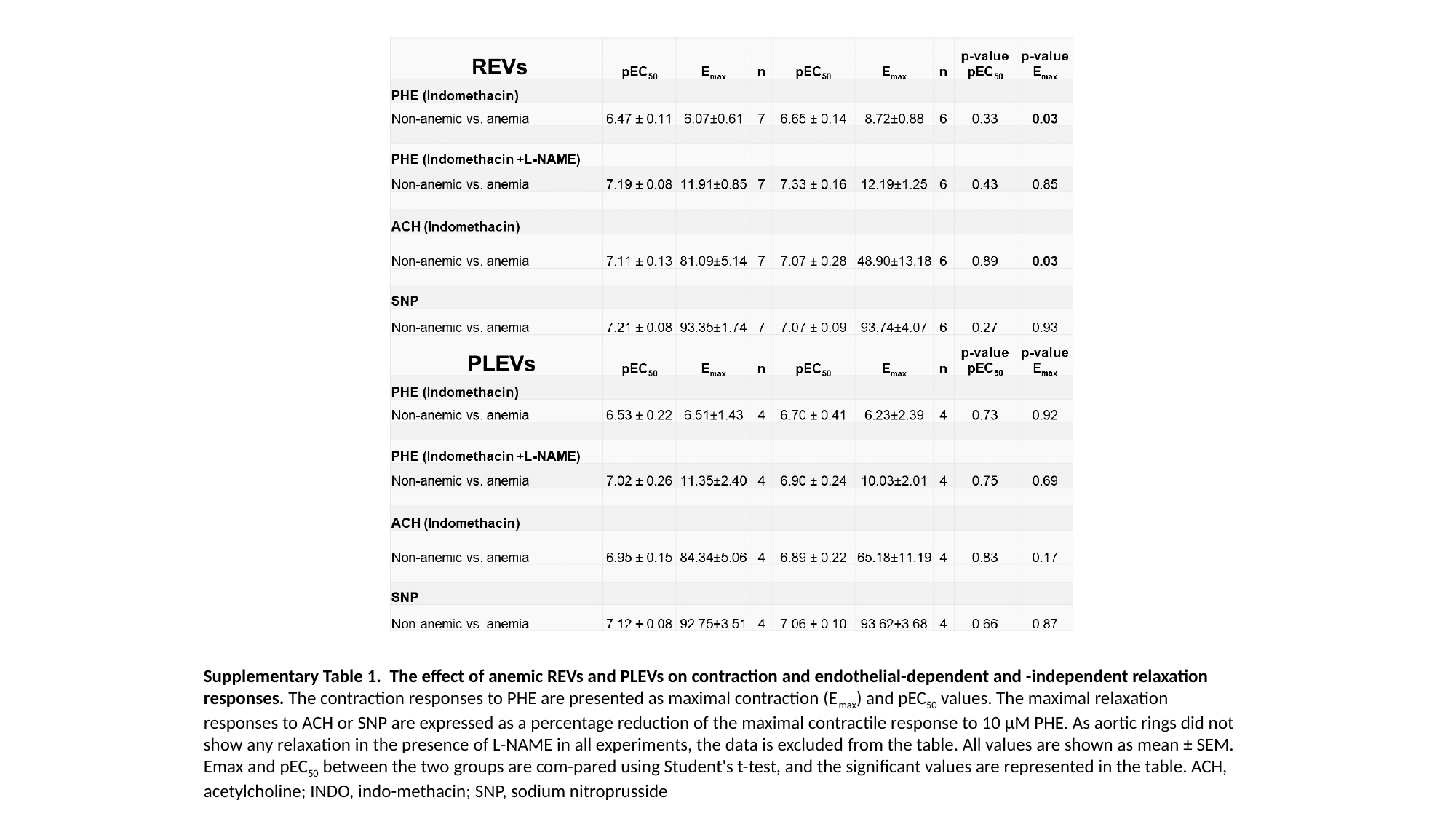

Supplementary Table 1. The effect of anemic REVs and PLEVs on contraction and endothelial-dependent and -independent relaxation responses. The contraction responses to PHE are presented as maximal contraction (E_max_) and pEC_50_ values. The maximal relaxation responses to ACH or SNP are expressed as a percentage reduction of the maximal contractile response to 10 µM PHE. As aortic rings did not show any relaxation in the presence of L-NAME in all experiments, the data is excluded from the table. All values are shown as mean ± SEM. Emax and pEC_50_ between the two groups are com-pared using Student's t-test, and the significant values are represented in the table. ACH, acetylcholine; INDO, indo-methacin; SNP, sodium nitroprusside

### Slide 7
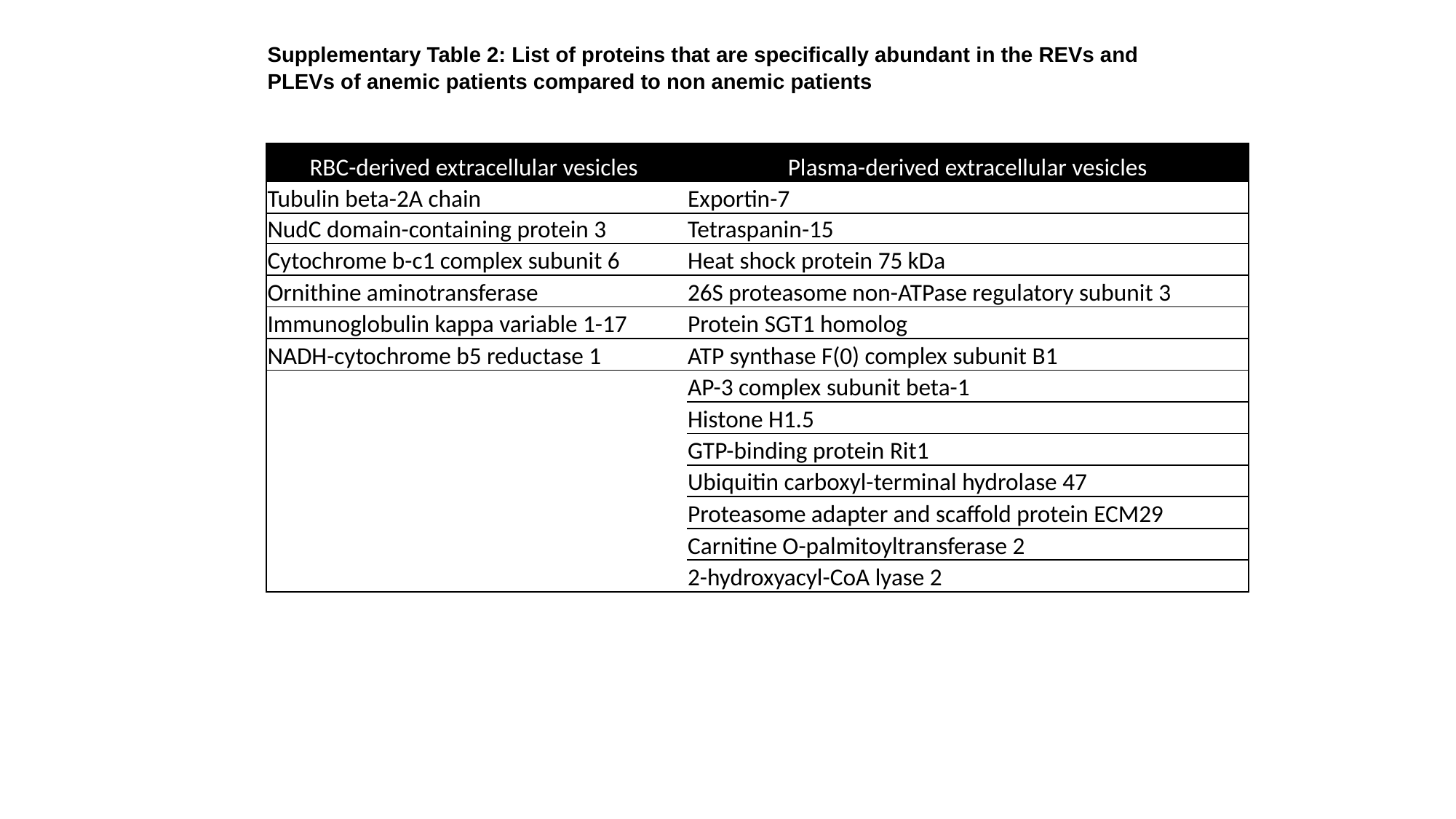

Supplementary Table 2: List of proteins that are specifically abundant in the REVs and PLEVs of anemic patients compared to non anemic patients
| RBC-derived extracellular vesicles | Plasma-derived extracellular vesicles |
| --- | --- |
| Tubulin beta-2A chain | Exportin-7 |
| NudC domain-containing protein 3 | Tetraspanin-15 |
| Cytochrome b-c1 complex subunit 6 | Heat shock protein 75 kDa |
| Ornithine aminotransferase | 26S proteasome non-ATPase regulatory subunit 3 |
| Immunoglobulin kappa variable 1-17 | Protein SGT1 homolog |
| NADH-cytochrome b5 reductase 1 | ATP synthase F(0) complex subunit B1 |
| | AP-3 complex subunit beta-1 |
| | Histone H1.5 |
| | GTP-binding protein Rit1 |
| | Ubiquitin carboxyl-terminal hydrolase 47 |
| | Proteasome adapter and scaffold protein ECM29 |
| | Carnitine O-palmitoyltransferase 2 |
| | 2-hydroxyacyl-CoA lyase 2 |

### Slide 8
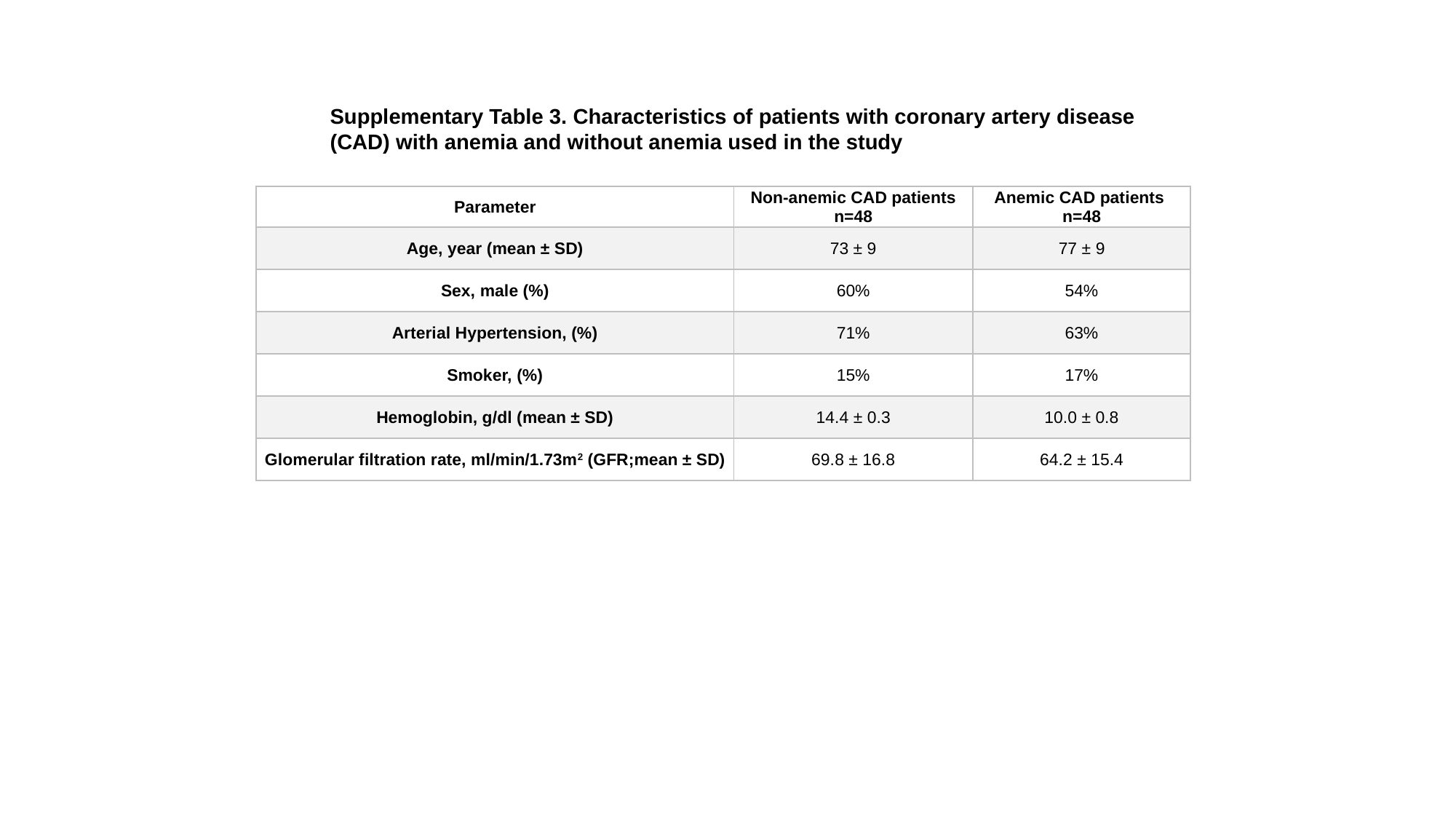

Supplementary Table 3. Characteristics of patients with coronary artery disease (CAD) with anemia and without anemia used in the study
| Parameter | Non-anemic CAD patients n=48 | Anemic CAD patients n=48 |
| --- | --- | --- |
| Age, year (mean ± SD) | 73 ± 9 | 77 ± 9 |
| Sex, male (%) | 60% | 54% |
| Arterial Hypertension, (%) | 71% | 63% |
| Smoker, (%) | 15% | 17% |
| Hemoglobin, g/dl (mean ± SD) | 14.4 ± 0.3 | 10.0 ± 0.8 |
| Glomerular filtration rate, ml/min/1.73m^2^ (GFR;mean ± SD) | 69.8 ± 16.8 | 64.2 ± 15.4 |
